# Individual bumble bees have small, unique, and persistent foraging repertoires: implications for disease transmission

**DOI:** 10.1101/2025.10.30.685600

**Authors:** Tallisker L.H. Weiss, Wee Hao Ng, Stephen P. Ellner, Christopher R. Myers, Scott H. McArt

## Abstract

Generalist species forage on a variety of resources, but individual members may specialize on a smaller subset of those resources. Previous attempts to incorporate intraspecific resource specialization in models of disease spread by pollinators have assumed specialization on one flower species, completely partitioning the plant-pollinator network, and very gradual changes in specialization, over days or weeks. These assumptions have not been demonstrated empirically but are critical to model conclusions. Here, we used palynology from sequential pollen loads of individual common eastern bumble bees (*Bombus impatiens*) to explore their foraging preferences and how they change over time. We found individuals in a bumble bee colony exhibited unique repertoires of two to three flower species while the full-colony repertoire was up to three times larger and varied by time of year. Floral constancy of individual bees lasted roughly ten days before turning over. We developed a disease transmission model parametrized with these empirical results, finding that R_0_ decreased with each additional flower species in a repertoire, which reduces the separation between theorized disease transmission subnetworks. Our results provide novel insight into the foraging behaviors of social bumble bees and improves understanding of how disease transmission occurs in complex plant-pollinator networks.

## Introduction

Pollinators such as bees are essential to natural and managed systems, as 87% of flowering plants require pollination (Ollerton et al., 2011) and one-third of our food is dependent on pollinating insects (Klein et al. 2007). Yet managed and wild pollinators are currently experiencing declines worldwide, in part due to pathogens (Cameron et al., 2011; Meeus 2011; Goulson et al., 2015). Pollinators transmit pathogens within and among colonies/nests and in community-wide plant-pollinator networks (Page et al. 2025). While solitary species are well-known to harbor pathogens (Evison et al. 2012; Ravoet et al. 2014; Graystock et al. 2020), generalist social species often have a disproportionate impact on pathogen prevalence in the full community. This is due to the fact that generalist social species live in communal groups where intracolony transmission is facilitated (Schmid-Hempel & Schimid-Hempel 1993, Schmid-Hempel 1995) and they are often numerically dominant in communities (Graystock et al. 2020; Cohen et al. 2021, Tiritelli et al., 2024). Both of these factors are conducive to pathogen spillover (Fürst et al. 2014; Müller et al. 2019; Manley et al. 2019; Piot et al. 2022).

Pollinators visit a wide range of plant species where pathogen transmission can potentially occur at flowers (McArt et al. 2014; Graystock et al. 2016; Figueroa et al. 2019). Because of this, plant-pollinator contact networks are often used to explore how transmission of pollinator pathogens occurs in the environment, with each node representing a species (Truitt et al., 2019; Figueroa et al. 2020; Manley et al. 2023). However, within generalist species, individuals may specialize on different subsets of the floral resources available, which can lead to differences between individual- and species-level contact networks (Araújo et al., 2011; Dupont et al., 2011). Individual specialization within generalist species is common (Bolnick et al., 2003, Bolnick et al., 2007; Araújo et al., 2011; Matich et al., 2011) and has been observed in a wide range of taxa, including sea otters (Estes et al., 2003; Woo et al., 2008; Newsome et al., 2009), loggerhead sea turtles (Vander Zanden et al., 2010), sparrows (Maldonado et al., 2019), arctic foxes (Angerbjörn et al., 1994), frugivorous bats (Kerches-Rogeri et al., 2020), ants (Rissing et al., 1981), and bees (Heinrich et al. 1976; Grüter et al., 2011). Thus, individual specialization has the potential to change transmission dynamics in individual-vs. species-level contact networks.

Previous theoretical work has found that mathematical models that ignore individual specialization can underpredict the persistence of many bee diseases, while either over- or under-predicting steady-state disease prevalence (Ellner et al., 2020). This is because individual specialization partitions the contact network into subnetworks, which limits pathogen spread via partitioning, but can also facilitate greater-than-average transmission within some subnetworks containing species that are especially conducive to transmission. However, several parameters from these models have not been assessed empirically. Specifically, while floral constancy is typically defined as a bee collecting pollen from a single flower species during a single foraging bout (Waser 1986), recent work has shown that individual bumble bees collect pollen from multiple species during individual foraging bouts (Gervais et al. 2020; Martinez-Bauer et al. 2021; Yourstone et al., 2023). Yet little is currently known regarding whether individuals develop unique foraging repertoires, how individual foraging diversity compares to full-colony foraging diversity, and how quickly individuals shift their foraging preferences. These parameters are all critical for determining whether or not a disease will persist on a multispecies plant-pollinator transmission network, and if so, what the endemic prevalence will be in each species (Ellner et al., 2020).

In this study, we collected and analyzed the pollen loads of individually identified bumble bees over multiple foraging bouts to address four major questions regarding individual specialization: 1) Are individuals constant to one or multiple species during a foraging bout?, 2) Do individuals visit unique repertoires of flowers over multiple foraging bouts? 3) How does individual foraging diversity compare to full-colony foraging diversity? and 4) What is the time frame over which individual foragers change their floral preferences? Based on these empirical results, we parameterized and tested a new model of disease transmission with the objective of predicting the influence a dynamic multi-species repertoire has on disease persistence in plant-pollinator communities with subnetworks.

## Methods

### Field experiment

Three trials were completed over the course of two summers in Ithaca, New York. All of the trials used commercially sourced *B. impatiens* colonies. The first trial was conducted during the summer of 2022 at Dyce Laboratory (42.4650474, −76.445121). For this trial, we set up one colony sourced from Biobest and tagged 250 worker bees each with a unique number so that we could identify which worker each pollen sample was collected from (BetterBee queen tags affixed with Loctite super glue). Pollen samples were collected from foraging bees from July 21^st^ to 25^th^ (see pollen collection protocols below). The second and third trials were conducted in the summer of 2024 at the Liddell Field Station (42.460788, −76.44394). Two new research colonies sourced from Koppert were used for each trial, and ~150 worker bees tagged in each colony (BetterBee queen tags affixed with Meyer Bees Queen Marking Number Glue). To distinguish between bees by their colonies of origin, bees from one colony were marked with red tags while those from the other was marked with blue tags. Trial 2 pollen samples were collected from July 22^nd^ to August 4^th^, and Trial 3 pollen samples from August 10^th^ to 23^rd^. Data collection occurred between 8 am and 5 pm daily for all three Trials.

Each of the two colonies was connected to a tunnel with an entrance/exit outside the building, allowing the bees to forage naturally. One of the setups is shown in Figure 1a. Each tunnel was monitored with a camera connected to a Raspberry Pi computer with a pre-trained detection algorithm (Figure 1c), and equipped with a gate at each end. When a tagged bee walked through the tunnel when entering the hive, its picture was taken (Figure 1b), and an alarm buzzed. If no pollen was found, we allowed the bee to enter the hive (Figure 1d). If the bee did have pollen on it, we closed the gates to stop the bee from leaving the tunnel (Figure 1e), pulled the bee out of the tunnel and placed it in a test tube. The bee was then moved into the freezer for a few minutes until it was calm enough for us to remove the pollen. Note that some pollen samples may have been missed when a tagged bee returned to the hive outside of the data collection time or when the observer was preoccupied with removing the pollen from another bee. The tools that were used to remove the pollen were sanitized between bees. Once the pollen sample was collected, the bee was returned to the hive. The identity of the bee was recorded in addition to which colony it entered with the pollen, to detect potential drift between colonies (bees leaving one colony and returning to the other).

**Figure 1:**
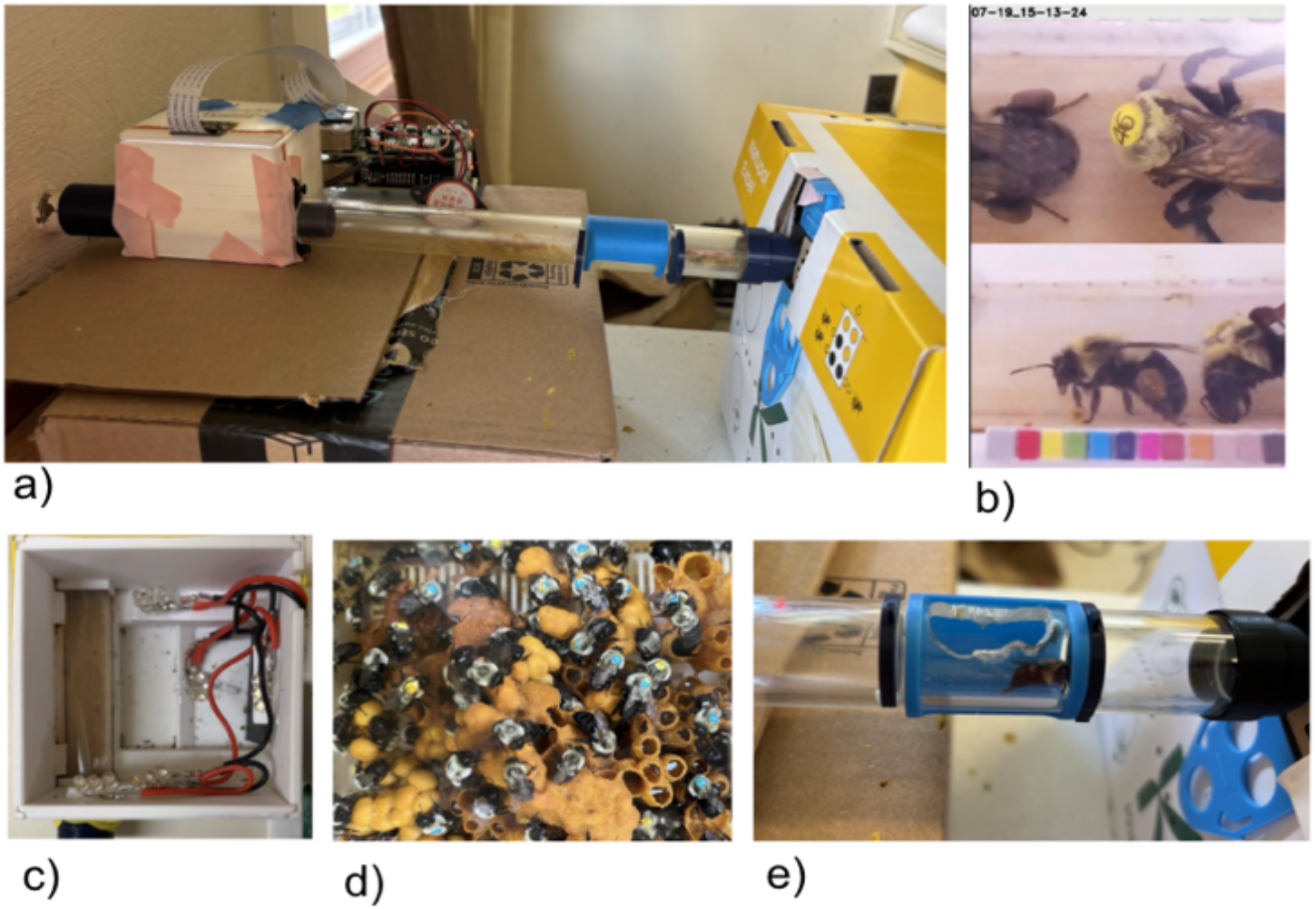
Experimental set-up for a bumblebee colony used for pollen sampling. **a:** Image capturing device attached to modular tunnel connecting the bumble bee colony (right) to the outside (left), to monitor bee entry/exit. **b**: Sample images of the top and side of a bee entering the hive. **c**: Inside the Raspberry Pi monitoring device. The camera is on the left side and is pointed at the transparent plastic tunnel surrounding the wooden board. Lights were used to maintain consistent illumination across all images to determine whether the bee was entering or exiting the tunnel. **d:** Blue and yellow tags on the bumble bees. **e**: The gates closed, and the trap door opened to allow the bee to be removed from the tunnel to be chilled and its pollen collected.

### Palynological assessment

Once pollen was collected, it was stored in a freezer before being screened using the methods described in McArt et al. (2017). Briefly, to prepare the slides, 400μL of DI water was added to the sample. It was then vortexed for 15 seconds before being placed in the centrifuge at 10,000 rpm for three minutes. The excess water was removed via micropipette, and 200 μL of 95% ethanol was added. The microtubes were vortexed for 10 seconds. Next, 10 μL of the sample was micro-pipetted onto a clean glass slide, which was mixed with 40 μL of Calberla’s solution. Calberla’s solution consisted of 5 mL glycerol, 10 mL 95% ethanol, 15 mL distilled water, and two drops of saturated aqueous solution of basic fuchsin. After 20 minutes, a cover slip was added and sealed to the slide the next day with clear nail polish.

On each slide, 300 pollen grains were counted under a microscope and categorized by morphotype (McArt et al., 2017; Urban-Mead et al., 2023). These morphotypes were not identified to species but were categorized based on their shape and size. Identifying marks such as the presence of spines, were also used to organize the groups. While a pollen type of less than 3% is usually considered to be the threshold for contamination, we did not remove those counts (except in Figure 2) as we chose methods of analysis that were not sensitive to such small numbers.

**Figure 2:**
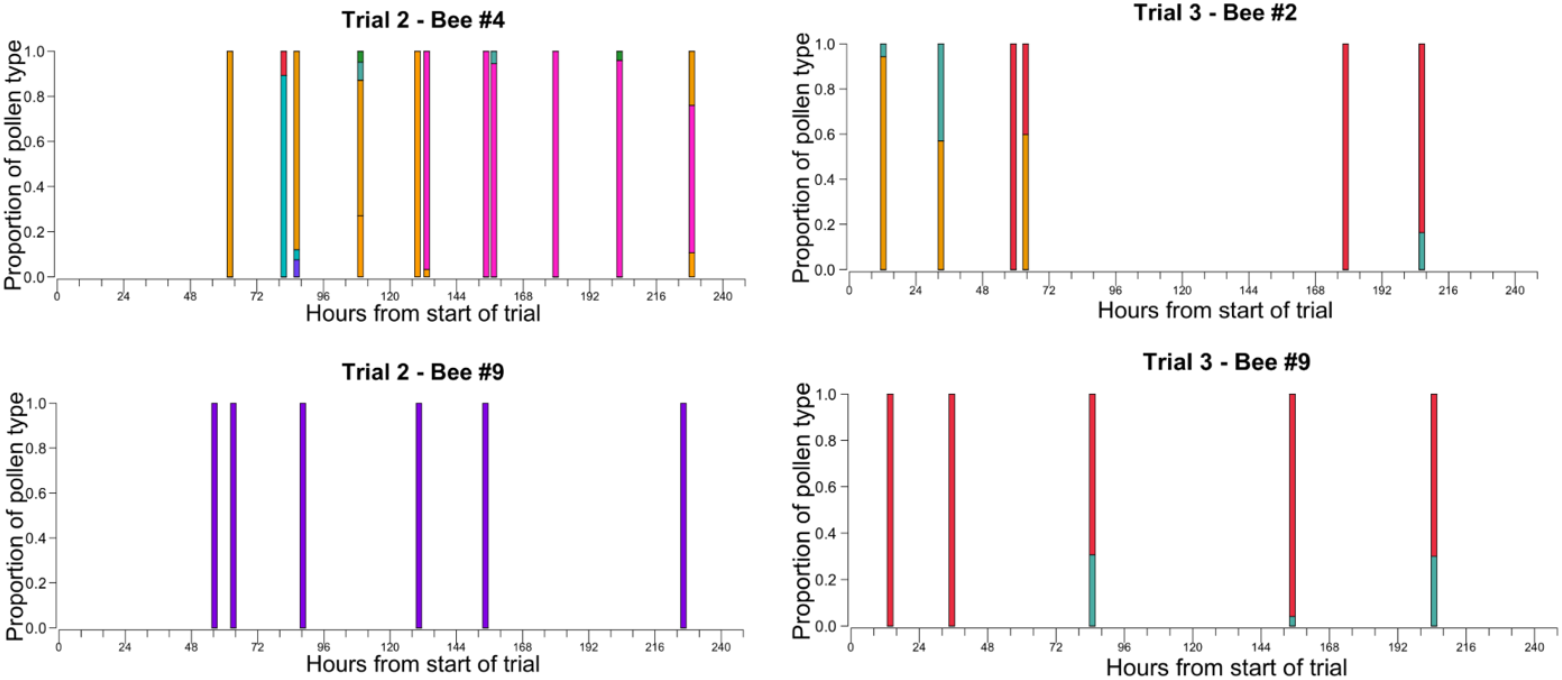
Four representative examples of sequential pollen foraging by individual bumble bee workers (*B. impatiens*) over 10 days. Each bee has a unique pattern with one to three main pollen types, that may change over time. The y-axis is the proportion of each pollen type, identified via microscopy, and the x-axis is the time of pollen collection in hours starting from time 0, which is the beginning of the respective trials. Each color represents a different pollen morphotype. For this figure, we used a 3% threshold as contamination (Louveaux et al. 1978); pollen designated as contamination was removed from the figures for visual clarity. See supplement Figure S2 for bar plots from all of the bees with sequential pollen samples.

All pollen samples for Trial 1 were screened, including those from bees with only one pollen sample collected (73 samples from 47 bees, average of 1.6 samples per bee); however, since we were interested in assessing floral constancy of individual bees, the analysis focused on the bees with multiple samples. Given the larger sample size of pollen, for Trials 2 and 3 we chose to focus on the bees that provided the most information about floral constancy. For Trial 2, a subset of the bees was screened (80 samples from 11 bees, average of 7.3 samples per bee). We chose bees that had multiple samples across both long- and short-time scales. Trial 3 had fewer samples than Trial 2, so all of the samples from bees that had three or more pollen samples collected from them were screened (47 samples, 12 bees, average of 3.9 samples per bee).

### Statistical Analysis

As a metric to characterize the compositional distribution of a typical pollen sample, we calculated the effective number of pollen types in each sample using the Simpson diversity index (Simpson, 1949) or equivalently, the Hill number with q = 2 (Hill, 1973). We chose the Simpson index, first because it is relatively insensitive to pollen contamination, and second because its values often agreed with what an effective number of pollen types intuitively ought to be. For instance, consider a pollen sample containing *n* pollen types. If a bee split its time evenly between all *n* types during a bout so they were present in nearly equal proportions, then Simpson index would be close to *n*, whereas if one pollen type dominated the sample, then Simpson index would become much smaller, approaching one in the limit of the dominant type reaching 100%.

While a single pollen sample represents one foraging bout, because a bee may choose to focus on different pollen types over the course of multiple bouts, a single pollen sample may not fully represent her current repertoire. To get a better sense of what the repertoire looks like, we calculated the effective number of pollen types across multi-day windows instead of a single pollen sample. More specifically, we pooled all pollen samples from the same bee within a moving window of *D* days and then calculated the Simpson diversity. We repeated this for different choices of *D*: shorter windows better reflect the repertoire at any time point if there is change in pollen repertoire over time, but may not represent the full repertoire of the individual bee. We focused on *D* from one to four days for Trial 1 and *D* from one to five days for Trials 2 and 3. For each value of *D*, the more pollen samples in a window, the more the effective number of pollen types we are likely to find, so we plotted the effective number of types as a function of the number of samples, analogous to a species accumulation curve; with increasing number of samples, the curve should eventually saturate at the true effective repertoire size. As reference, for each number of samples we also pooled that many randomly chosen pollen samples from different bees across the entire trial duration, and calculated the effective number of types; we repeated this 50 times and calculated the median. This reference is representative of the colony-level (as opposed to individual-level) effective pollen diet breadth during the trial.

To determine if the pollen composition of samples collected differed among bees, non-metric multidimensional scaling (NMDS) was used for ordination and visualization (Kruskal, 1964), while permutational multivariate analysis of variance (PERMANOVA) was used to test for differences in the composition (Anderson, 2001). A weighted bipartite network comprising individual bees and pollen types as nodes was created for additional visualization. Weighted Jaccard distance was used as the dissimilarity metric because unlike binary Jaccard distance, it takes into account the relative abundance of the pollen types and is also less sensitive to pollen contamination. Note that for Trial 1, we only used bees with at least 3 pollen samples for the NMDS and PERMANOVA analyses. The R package **vegan** (Oksanen et al., 2022) was used for these calculations.

As yet another way to assess whether each bee had a much smaller repertoire than the colony as a whole, and also whether the repertoire changed with time across each trial, we calculated the weighted Jaccard distance between every pair of pollen samples. This was used to assess whether there was a difference in similarity between pairs of pollen samples collected from the same bee (henceforth “same-bee pairs”) and from two separate bees (henceforth “different-bee pairs”). Additionally, it was also used to determine if pairs of pollen samples become more different as the time separations between the pairs increase, indicating a change in the pollen selection choices over time for an individual bee. We first fitted a generalized linear mixed models (glmm) with the response variable being the weighted Jaccard similarity (1 - Jaccard distance). We used the ordered beta family as it can take exact zeros and ones. We included a binary variable for whether a pair was same- or different-bee, the time separation of a pair, and their interactions as fixed effects. To account for pseudoreplication, we used bee ID pairs (which would be the same bee ID twice for “same-bee pairs”) as the random effects grouping variable of the pollen pairs. The random effects included random intercepts and slopes, but structured in a way such that same- and different-bee pairs could draw random intercepts and slopes from different normal distributions. We also allowed the dispersion of the ordered beta distribution to vary between same- and different-bee pairs. Because the interactions between the binary variable and time separation were not significant (Trial 1: z=1.3, p=0.2; Trial 2: z=-0.42, p=0.67; Trial 3: z=1.3, p=0.20), by the principle of marginality we refitted the model without the interaction to assess the main effects of the binary predictor and time separation. The GLMMs were fitted using the R package glmmTMB (Brooks et al., 2017). One concern is that the similarities are not independent, in that modifying one pollen sample simultaneously changes the similarities of all pollen pairs containing that sample, so the model may overestimate the precision of the fit. To confirm the significance of the positive results, we performed permutation tests; see supplement section S2. Finally, because some bees were observed to drift between the two colonies during Trials 2 and 3, we tested whether this could have affected the Jaccard similarity of the same-bee pollen pairs, but did not find any significant effects (see supplement Section S1 for details).

### Model for disease transmission in a plant-pollinator network

Full model details are provided in Supplement section S3. Briefly, we constructed three differential equation models, each representing a different assumption about the foraging repertoire of a bee. For each model, we generated 2,500 random sets of parameters, and for each parameter set compared the values of the basic reproduction number R_0_ when a bee sticks to a single repertoire for life (“slow switching”), versus when a bee rapidly switches between repertoires and effectively becomes a generalist (“fast switching”). Because R_0_ is a measure of the ability for a disease to spread and persist, the difference in R_0_ between the two slow- and fast-switching limits indicated the how much we would under- or overestimate disease persistence had we neglected repertoire constancy. By repeating the calculations for all three models, we could assess how this consequence depended on our assumptions about the repertoire.

The first model (“single major”) is the Ellner et al. (2020) model where a bee only forages on one flower species at a time. In contrast, both the second and third models assume repertoires of effective size two (as represented by the Simpson index), but implement them in two different limits: the second model (“double major”) assumes each bee forages equally from two flower species, while the third model (“random sampling”) assumes that each bee has a single specialty but occasionally performs “random sampling” trips where she visits all other species in the community such that a Simpson index of two is still achieved; see supplement S3.

Since our goal was to assess how the consequence of neglecting constancy depended on the model assumptions, to facilitate comparison between the three models, we wanted all three models to reduce to the same “generalist model” in the fast-switching limit. Hence, the parameter values for our simulation studies were drawn at random, but constrained such that they all produced the same species-level foraging proportions. All three models therefore have the same R_0_ for fast switching but different R_0_ values for slow switching due to the different assumptions about repertoires.

## Results

In total, we screened 200 total pollen loads from 70 bees (Table 1): 47 bees in Trial 1, 11 bees in Trial 2, and 12 bees in Trial 3. Pollen samples from a single bee were typically spread out over multiple days, and frequently there were multiple samples from individual days (see Figure 2).

**Table 1:** Summary of when each trial was conducted, the number of worker bees (*B. impatiens*) tagged, and pollen samples collected from and screened in each trial. Screening refers to using microscopy to identify and count the morphotypes found in the pollen sample. Not all samples were screened in Trial 2 and 3. We screened all pollen sampled from Trial 1, while the screening for Trials 2 and 3 focused on bees that were repeatedly sampled.

|  | Date of Trial | Number of bees initially tagged | Total number of pollen samples collected | Total number of bees with the pollen removed | Number of pollen samples screened | Number of bees whose pollen was screened |
| --- | --- | --- | --- | --- | --- | --- |
| Trial 1 | July 21 – July 25, 2022 | 250 | 73 | 47 | 73 | 47 |
| Trial 2 | July 22 – August 4, 2024 | 150 | 429 | 201 | 80 | 11 |
| Trial 3 | August 10 - August 23, 2024 | 150 | 105 | 72 | 47 | 12 |
| Total |  | 550 | 607 | 320 | 200 | 70 |

Figure 2 shows four representative examples of sequential pollen foraging by individual bees over 10 days. The average time between consecutive pollen foraging bouts was 5 days, and a maximum of 11 days during which a bee was observed. Figure 3a shows that the majority of the individual pollen samples are dominated by one pollen type, this is especially the case for Trial 2. This pattern is further confirmed by the effective number of pollen types having a median close to one, although there appears to be more spread in Trial 3 (Figure 3b).

**Figure 3:**
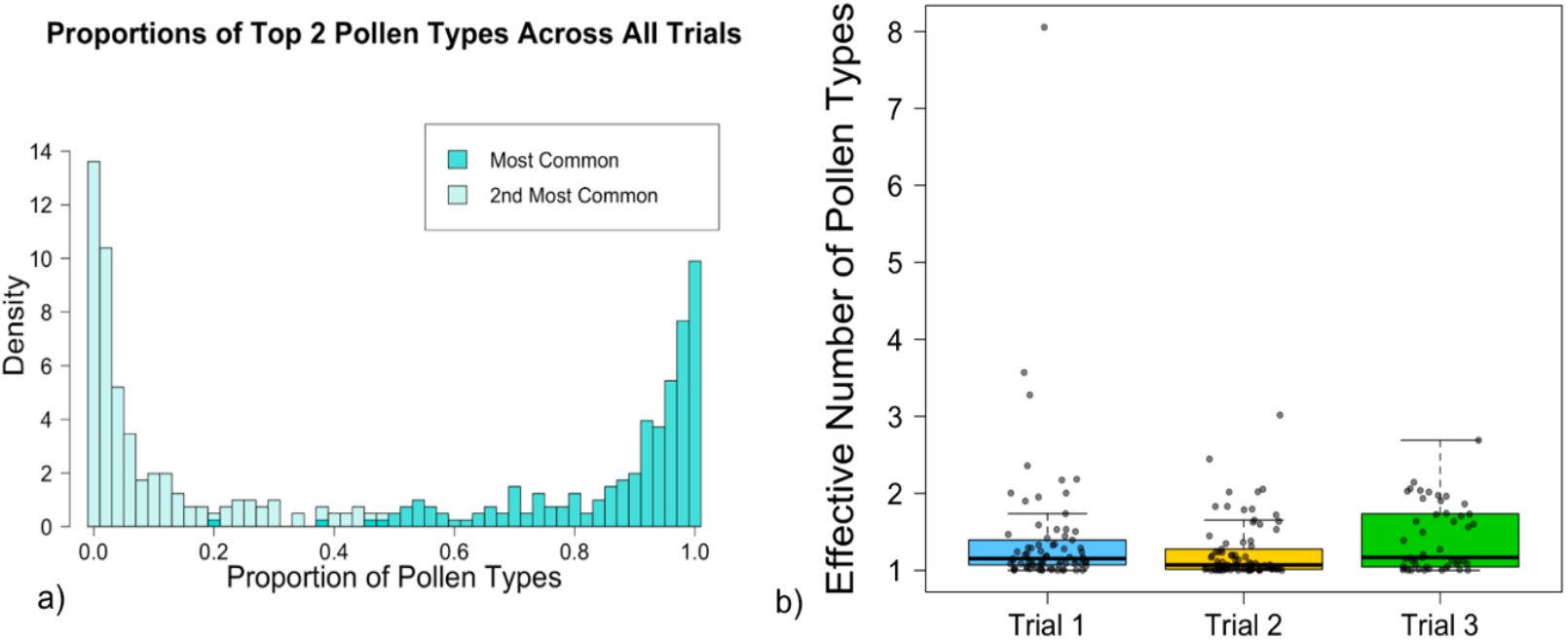
Most pollen samples collected from individual bumble bee (*B. impatiens*) workers are dominated by a single pollen morphotype. **a:** For each pollen sample, we calculate the proportions of the most common and second most common pollen types. Here, we show histograms of these proportions for pollen samples across all three trials; see supplement for the histograms separated by trial. **b:** The effective number of pollen types based on the Simpson diversity index. Each point represents one pollen sample.

The NMDS plots for the three trials (Figure 4) show clear separation between each individual bee’s foraging preferences which was confirmed using PERMANOVA (p=0.001 for Trial 1, p=0.001 for Trial 2, p = 0.001 for Trial 3); this indicates that each bee has its own foraging repertoire. Note that for Trial 3, many samples were compressed along a line (near NMDS2 = 0), probably because most of these samples contained a substantial proportion of pollen types 30 and/or 31.

**Figure 4:**
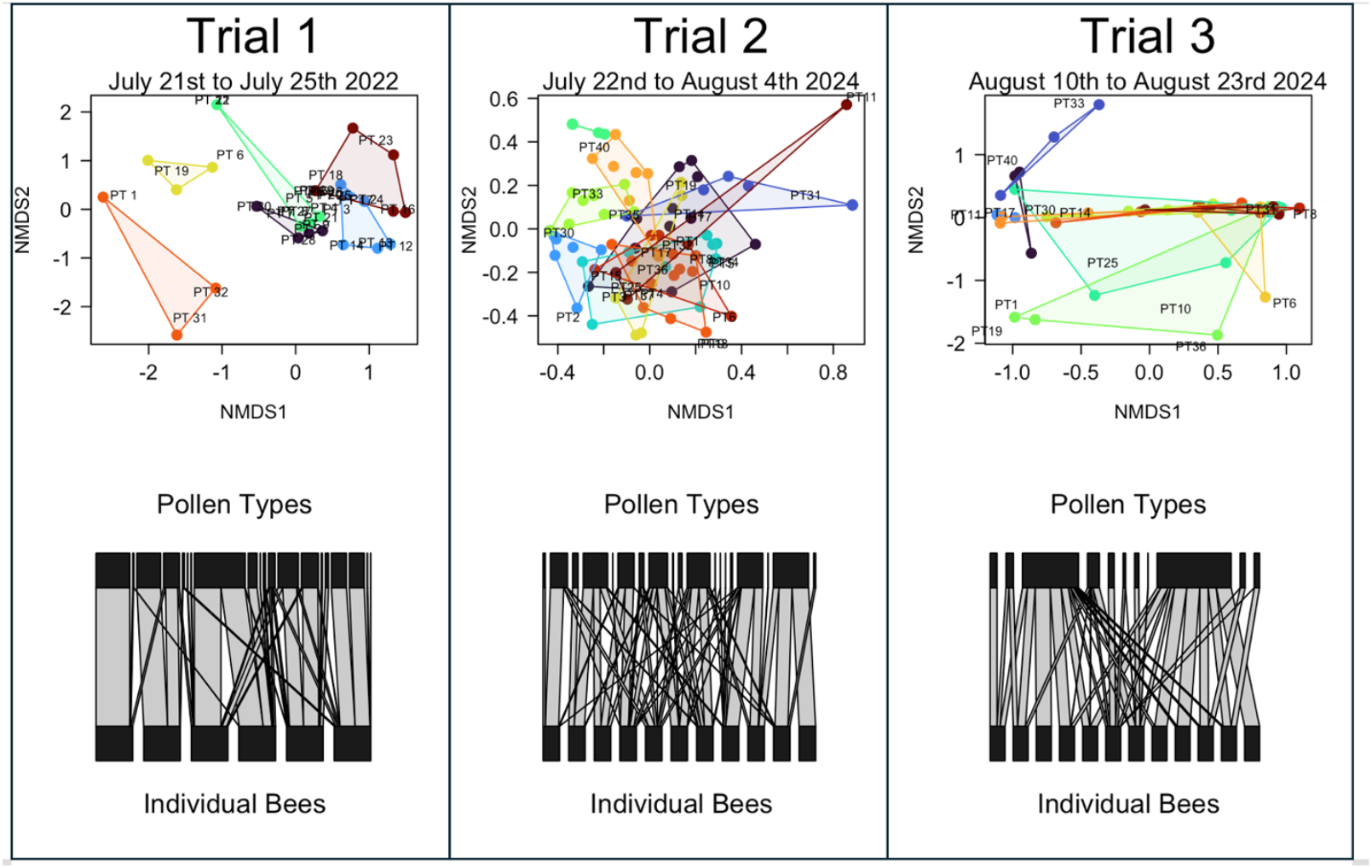
Pollen samples obtained from individual bumble bee (*B. impatiens*) workers show that bees have distinct repertoires that are much smaller subsets of the full-colony repertoire. Each panel shows one trial, with a non-metric multidimensional scaling (NMDS) ordination plot on top and a bipartite network plot on the bottom. The NMDS was created with the weighted Jaccard distance between pollen samples as the dissimilarity index. Each point is a pollen sample and each color represents an individual bee. Only bees with three or more consecutive pollen samples are included. The polygons are the convex hull of the pollen samples from a bee. “PT” refers to the pollen type (and associated number) identified, where a pure sample of that particular pollen type would be at the center of the label, so a pollen sample close to a label is mostly dominated by that pollen type. For example, in Trial 1, the orange triangle in the lower left corner shows that the bee completely switched the flower it collected pollen from three separate times between PT 1, PT 31, and PT 32. For the network analysis, the top black bars represent the different pollen types, while the bottom black bars represent individual bees. The grey bands connect individual bees to the pollen types that they collected, with wider bands indicating higher proportions. The width of a pollen type bar is the sum of the widths of the gray bands connected to it, so the more popular a pollen type, the wider the pollen type bar. For Trial 3, the two largest bars correspond to pollen types 30 and 31, which is consistent with the NMDS ordination plot where many points are also clustered around these two pollen types.

For a 5-day window, with an increasing number of pollen samples, the effective number of pollen types in the pollen samples from a single bee in Trials 1 and 2 appeared to asymptote towards two to three pollen types, which is much lower than that of the reference levels based on random samples from different bees (Figure 5). This indicates that each bee does not only have a single specialty, but the individual repertoire is still only a small subset of the colony-level diet breadth. For Trial 3, the individual and reference asymptotes are very similar (Figure 5), indicating the colony-level diet breadth was not substantially larger than the individual repertoires.

**Figure 5:**
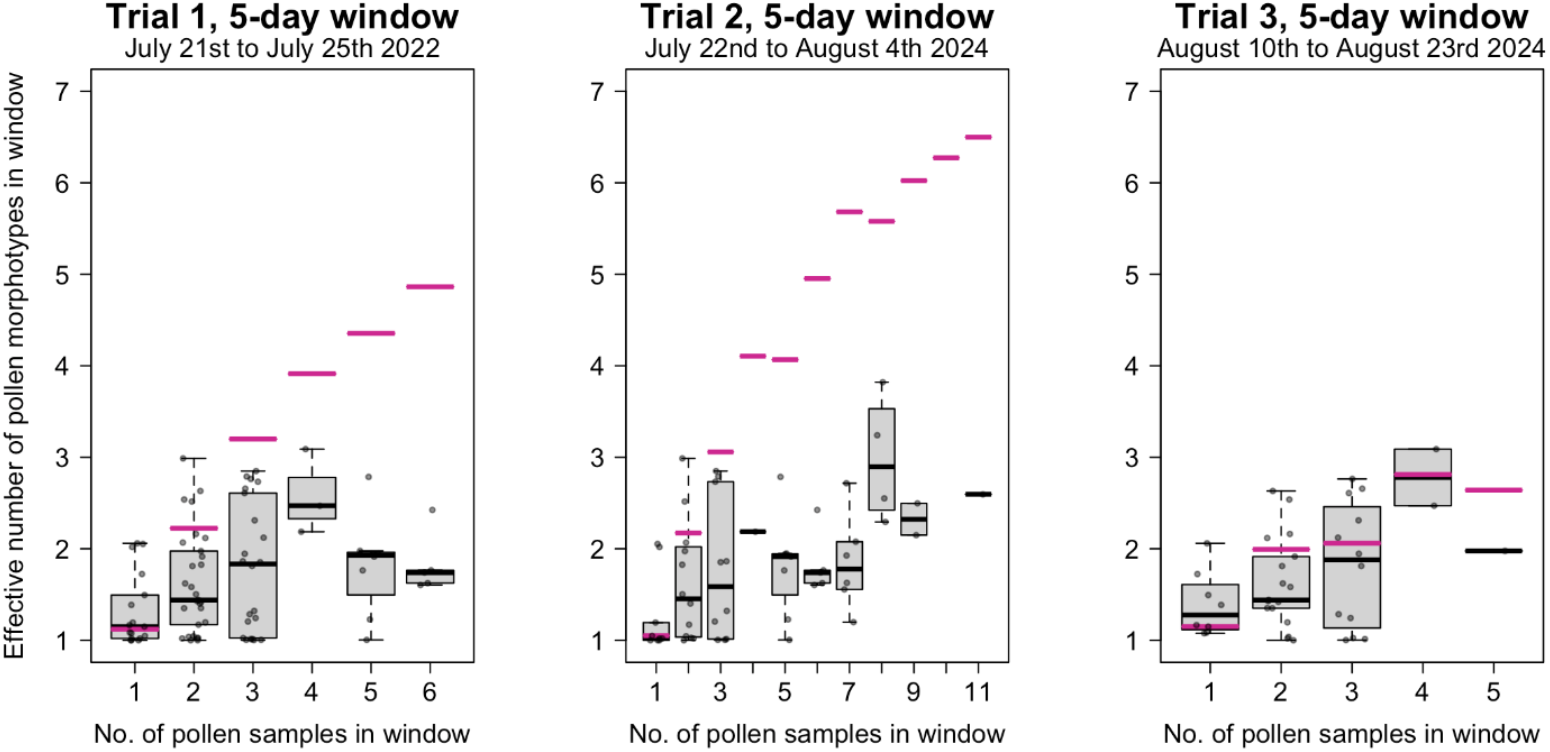
The repertoire of individual bumble bee (*B. impatiens*) workers over multiple foraging bouts comprises about two to three main pollen types (boxplots), regardless of colony-level foraging breadth (pink reference lines). Again we use Simpson’s diversity index as a measure of the effective number of pollen types. Each point corresponds to Simpson’s index of all pollen samples from the same bee pooled within a moving window of 5 days; see supplement S4 for other window lengths. Because Simpson’s index may depend on the number of pooled samples, each panel is analogous to a species accumulation curve. The pink lines were calculated using randomly selected pollen samples from different bees and hence represent the equivalent Simpson’s indices for the full colony’s repertoire.

For the analysis of Jaccard similarities, we found that in all three trials, pollen samples from the same bee (same-bee pairs) are less different than pairs of pollen samples from different bees (different-bee pairs) (Figure 6; Trial 1: z = 8.179, p_glmm_ < 0.001, p_perm_ < 0.001; Trial 2: z = 7.792, p_glmm_ < 0.001, p_perm_ < 0.001; Trial 3: z = 2.614, p_glmm_ < 0.001, p_perm_=0.002), indicating that individual bees had smaller repertoires than the colony-level diet breadth. For Trial 2, the similarity of both same- and different-bee pairs decreased over time (z = −3.191, p_glmm_ < 0.001, p_perm_ = 0.039). For Trial 1, we also saw a decrease in similarity with a very similar slope estimate, although it was only marginally significant, perhaps because of the shorter duration and smaller sample size (z = −2.1, p_glmm_=0.036, p_perm_=0.072). In contrast, for Trial 3, there was no significant effect of time separation on similarity (z = −0.043, p_glmm_ = 0.966). Finally, for Trial 2, at small time separations, the Jaccard similarity appeared to be bimodally distributed with a comparable number of pairs in both the high- and low-similarity peaks (Figure 6), which is consistent with each pollen sample being dominated by a single pollen type and a repertoire of two to three species.

**Figure 6:**
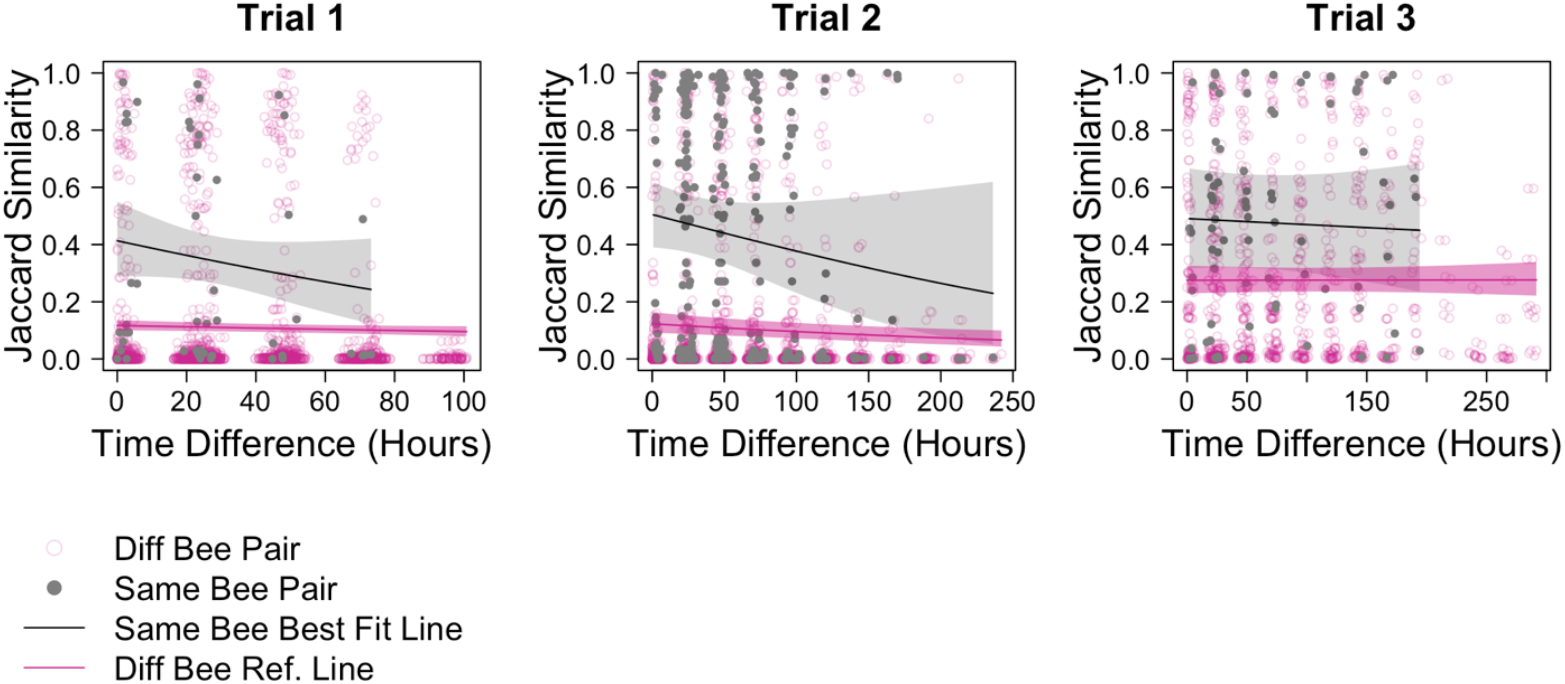
Individual bees exhibit floral preferences that change over time. Here, each point corresponds to a pair of pollen samples. Solid points (grey) are sample pairs from the same bee, while the hollow points (in light pink) are pairs from two different bees. The weighted Jaccard similarity (a metric of similarity in pollen composition) between two pollen samples is on the y-axis and the time difference (in hours) between pairs of samples on the x-axis. Black line (with 95% confidence shading) is the best fit for samples between the same bee. Pink line (with 95% confidence shading) is the best fit for pairs of samples from different bees. The points also appear to be bimodally distributed; this is consistent with Figure 2 showing that most individuals only focus on one species per foraging bout, so in each pair both samples were either dominated by the same type (very high similarity) or by two different types (very low similarity).

Based on these empirical results, especially our observation that individual bees appear to exhibit constancy not to a single pollen type but rather a repertoire of effective size between 2 and 3, we updated the Ellner et al. (2020) theoretical model for disease transmission in a plant-pollinator network as described in the methods, to re-evaluate how the consequence of ignoring floral constancy would change under different assumptions about a bee’s foraging repertoire. In Figures 7a-c for the single major, double major and random sampling models respectively, each point represents a randomly chosen set of parameter values, the horizontal position indicating the resulting value of R_0_ in the fast-switching limit (no constancy), and the vertical position the value of R_0_ in the slow-switching limit (lifelong repertoire constancy). Almost all the points lie above the 1:1 line (shown in dashed), indicating that in all three models, ignoring constancy would often cause one to underestimate R_0_ and hence disease persistence. Nonetheless, the effects appear to be smaller when the repertoire size was raised from one to two. More specifically, the slow-switching R_0_ in the double major and random sampling models were only half as large as the single major model. Since the increase in R_0_ from fast-to slow switching in the single major model was due to the partitioning of the plant-pollinator network into single-plant-species subnetworks (Ellner et al. 2020), the reduced effects in the repertoire-size two models may be due to the increased repertoire size reducing the extent of the partitioning. Interestingly, the results found for double majoring and random sampling were almost identical despite being two very different ways of achieving a repertoire size of two, suggesting that the effects of floral constancy on disease transmission may depend more strongly on the repertoire size (as defined by the Simpson index) than on how that repertoire size was achieved.

**Figure 7:**
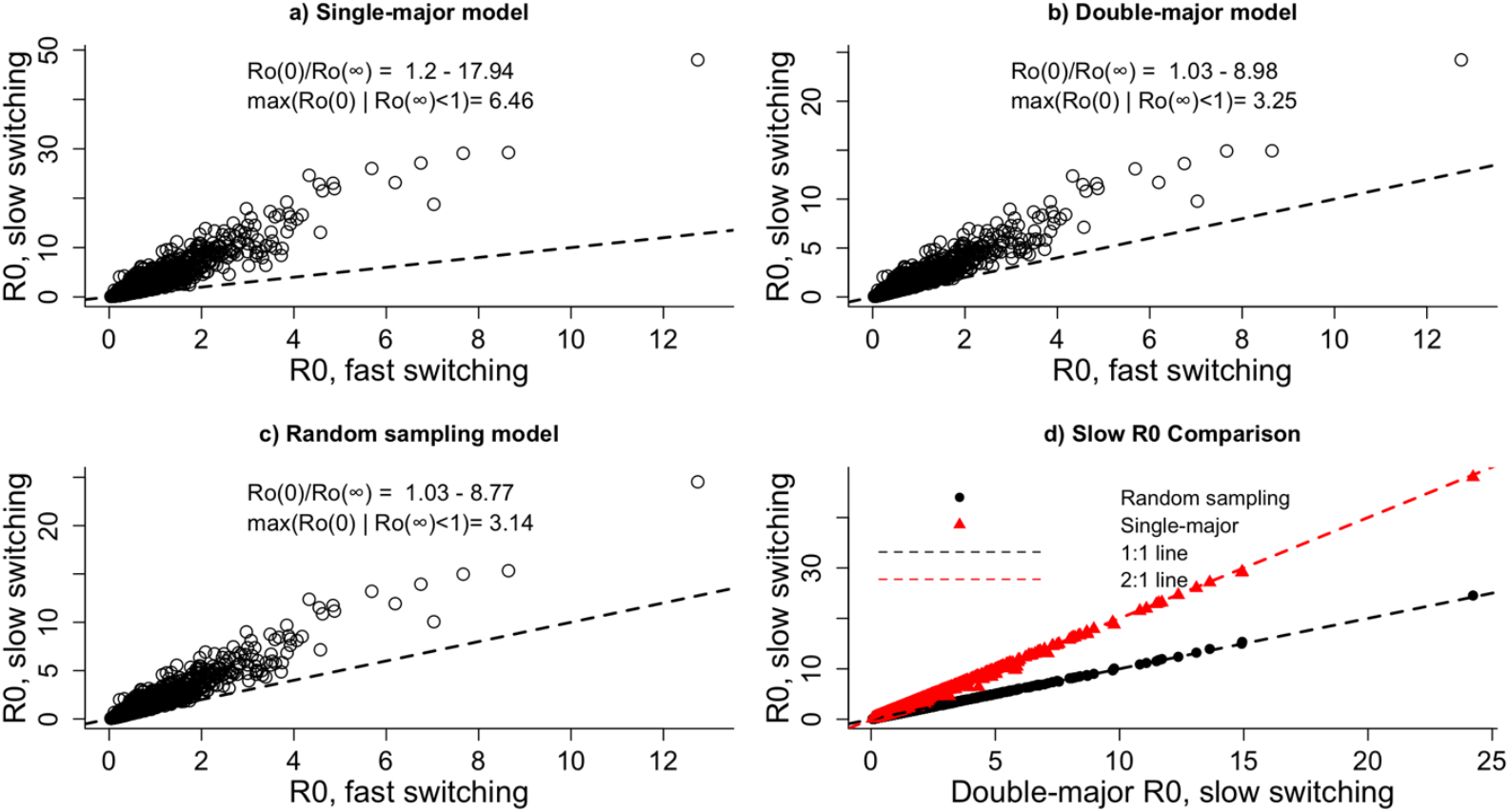
Ignoring floral constancy in foraging behavior leads one to underestimate bee disease persistence, but to a smaller extent when the repertoire size of a bee is increased from one flower to two. Here we compare three mathematical models of bee disease transmission that incorporate floral constancy but with different assumptions about an individual bee’s repertoire: (a) single major model from Ellner et al. (2020), repertoire size one; (b, c) double major and random sampling models, each repertoire size two. Each point indicates a random set of model parameters, the x-axis corresponds to the basic reproduction number R_0_(∞) in the fast-switching limit where bees frequently change repertoires (hence losing floral constancy), and the y-axis corresponds to R_0_(0) in the slow-switching limit where a bee sticks to the same repertoire throughout her lifespan (lifelong constancy). The dashed line is the one-to-one line; because most points fall above the line so R_0_(0) > R_0_(∞), this means that ignoring floral constancy would often lead one to underestimate disease persistence. However, (d) shows that R_0_(0) in the repertoire-size two models are generally lower than that of the repertoire size one model, so the degree of underestimation is less severe.

## Discussion

In this study, we used a unique experimental setup that combined palynology with an automated bee monitoring and alert system to assess the extent and duration of floral constancy in *B. impatiens* workers. Individual bees typically had a distinct foraging repertoire of around two to three flower species regardless of the number of species utilized by the colony as a whole, which can be much larger. Hence, while bumble bees differ from honey bees in that they do not stay constant to a single flower species (Grüter et al., 2011), they still exhibit floral constancy in a more generalized sense by specializing, at any time, on a relatively small number of species. These results are consistent with Heinrich’s original observations (Heinrich, 1976), where he observed that bumble bees workers often have a small repertoire consisting of a major and a minor species. Further, we used the empirical results from this study to update a disease transmission model that explicitly considers individual specialization (Ellner et al., 2020), finding that R_0_ decreased with each additional flower species in a repertoire. Thus, ignoring floral constancy in foraging behavior leads to an underestimate bee disease persistence in plant-pollinator networks.

While the use of palynology to assess an individual bee’s foraging breadth is not new, the automated system greatly improved monitoring efficiency of foraging bouts compared to previous studies of floral constancy based on mark-recapture (Yourstone et al. 2023). For example, for the bees that we analyzed in Trial 2, the number of samples collected from repeated recaptures of a single bee ranged from 3 to 11 (median 7.5), much larger than the 2 to 4 samples in Yourstone et al. (2023). The increased rate of recapture and pollen sampling made it possible to uncover novel aspects of their foraging behavior. Specifically, we were able to identify the composition of foraging repertoires of individual bees and how they changed over very small time scales (within a day) or larger time scales (multiple weeks). Repertoires were unique and largely consistent for individual bees (Figure 4). Based on the decrease in the Jaccard similarity between pollen pairs collected from the same bee over time (Figure 6), a typical bee changed its repertoire over an average duration of roughly ten days.

Because pollen pairs collected from different bees exhibited decay over time, the loss of constancy may be mostly driven by floral community turnover, and it is possible that repertoires could have persisted longer in a less dynamic floral community. Indeed, in Trial 3 where the Jaccard similarity between pollen pairs from different bees did not decay over time, neither did the similarity between pollen pairs from the same bees. Likewise, Heinrich (1976) observed in his study that switching of preferences was relatively rare over a bumble bee’s lifespan, and attributed it to the long flowering duration of plants in the study site compared to the lifespan of a worker bee. We also observed that in Trial 3, two flower species showed up prominently in the repertoires of almost all the bees that were screened (Figure 4). Given that Trial 3 occurred later in the season, this could reflect the dominance of a small number of late-season flower taxa such as goldenrods and knapweeds in the region. Altogether, our results emphasize the important role of the floral community in shaping the foraging behavior of bumble bees at the individual level.

We acknowledge that our experimental approach has two major limitations. First, while an advantage of palynology over field observational studies (e.g., Heinrich, 1976) is that we are less likely to miss parts of or entire foraging bouts and hence underestimate the number of species utilized by a single bee, it also means that we can only assess pollen foraging behavior. We do not know whether the same floral constancy behavior equally applies to nectar foraging. This is important if we want to understand bee disease transmission since both pollen and nectar foraging can contribute to indirect transmission. Second, we have been referring to pollen morphotypes and flower species interchangeably, when it is possible that there may be multiple flower species with pollen morphotypes that cannot be distinguished by visual appearance alone. This limitation is not unique to our work; in Yourstone et al. (2023) which used a neural network for pollen identification, the authors found it necessary to group the flower species into a much smaller number of pollen morphotypes because the species in each morphotype were hard to tell apart even by the neural network. However, from a disease modeling standpoint, this limitation may not be particularly important if flower species with morphologically similar pollen also had similar traits relevant to disease transmission, and so do not need to be differentiated.

Our findings have two main implications for the epidemiology of bee diseases. First, our observations that bumble bees exhibit floral constancy means that epidemiological models of disease transmission of bumble bee diseases would need to incorporate foraging constancy behavior, as in the Ellner et al. (2020) model. However, the fact that bumble bees are constant to a larger repertoire of two to three species unlike the single species as assumed in Ellner et al. (2020) also suggests that the effects may be less pronounced because the overlap in flowers species between repertoires reduces the extent to which the plant-pollinator networks are partitioned by the constancy behavior. Indeed, we found (Figure 7) that the extent to which ignoring constancy underpredicts R_0_ was less severe in the repertoire-size-two models (double major and random sampling) than in the size-one model (single major). Nonetheless, even in the size-two models, the effects can still be very strong, producing in some situations large (up to five-fold) differences in the value of R_0_ with and without floral constancy. Hence, ignoring individual specialization and its persistence can still lead to predictions of disease die-off when in fact there is robust disease persistence (R_0_ of 1 or much higher).

Second, floral community turnover may determine the importance of individual specialization on disease transmission by driving switching behavior. An important parameter in Ellner et al. (2020) is the rate at which an individual changes its preference relative to the disease timescale. For example, a bee that frequently switches repertoire would be no different from a generalist with no floral constancy as far as the disease dynamics are concerned. In Ellner et al. (2020), the floral community was assumed to be static, and the rate of preference change was assumed to be an inherent trait of the species. In contrast, our results suggest that floral community turnover may play at least a partial role in driving any repertoire change. Therefore, the dynamics of the floral community cannot be neglected when trying to model the effects of individual specialization on disease transmission. Because the rate of community turnover may vary across the season, this also means that the effects of floral constancy on disease transmission may also vary, depending on the disease timescale.

Given that several taxonomic groups other than bees also exhibit individual specialization (Bolnick et al., 2003), accurate prediction of epidemiological dynamics in these taxa may require incorporating such behavior into disease transmission models. As our work here have shown, details about individual behavior, such as the repertoire size or the duration over which a repertoire is maintained, may vary between taxa and may also depend on the community that the taxa interact with. Models have proven to be an essential tool for simulating disease transmission, yet in order to improve their accuracy, empirical data are needed to parameterize individual behavior for the specific taxon and system of interest. Hence, our work highlights the importance of continued experimental or observational studies of individual behavior in understanding the dynamics of disease transmission.

## Supporting information

Supplementary Figures and Sections

## Acknowledgements

Thanks to Dr. Tom Seeley for his help setting up the hives and for the use of the Liddell Field Station. We would like to thank David Sossa and Dr. Leah Valdes for their help in tagging the bees. Thank you to the members of the McArt Lab for their support and feedback.

## Author contributions

TW, SM, SE, WHN, and CM conceptualized the study. SE developed and analyzed the single and double majoring models. CM assisted with the analysis of the analysis of the bee identification and pollen collection data. WHN worked on the figures, contributed to the analysis and study design, and created the random sampling model. SE, SM, WHN, and CM all provided input on the study throughout and contributed to the writing of the manuscript. TW collected all of the pollen samples, tagged the bees, wrote the initial manuscript, worked on the figures, and assisted in the random sampling model.

