## Supplementary Figures and Sections for "Individual bumble bees have small, unique, and persistent foraging repertoires: implications for disease transmission"

### Supplement

**Figure S1**: Images of the pollen morphotypes and their associated numbers. All images were taken at the same magnification but cropped and zoomed in for increased visibility.

| | | | | |
|---|---|---|---|---|
| Pollen Type 1<br>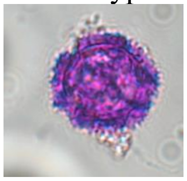 | Pollen Type 2<br>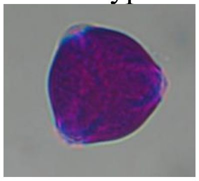 | Pollen Type 3<br>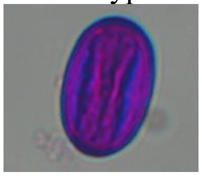 | Pollen Type 4<br>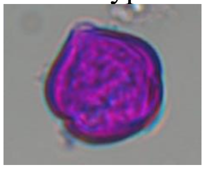 | Pollen Type 5<br>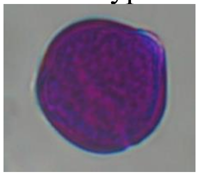 |
| Pollen Type 6<br>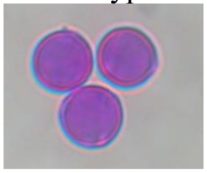 | Pollen Type 7<br>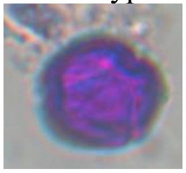 | Pollen Type 8<br>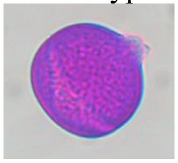 | Pollen Type 9<br>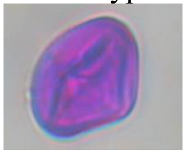 | Pollen Type 10<br>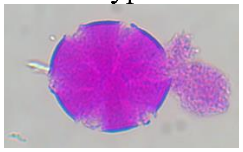 |
| Pollen Type 11<br>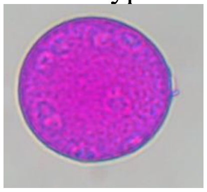 | Pollen Type 12<br>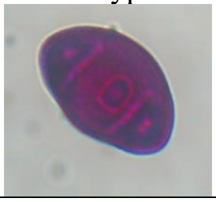 | Pollen Type 13<br>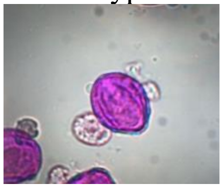 | Pollen Type 14<br>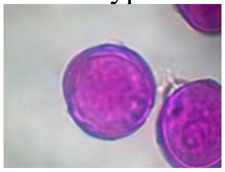 | Pollen Type 15<br>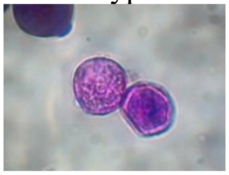 |
| Pollen Type 16<br>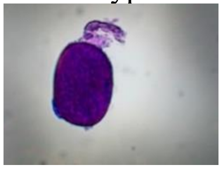 | Pollen Type 17<br>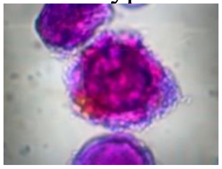 | Pollen Type 18<br>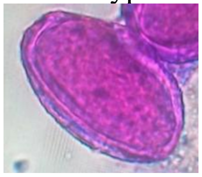 | Pollen Type 19<br>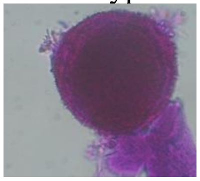 | Pollen Type 20<br>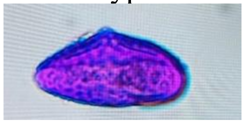 |
| Pollen Type 21<br>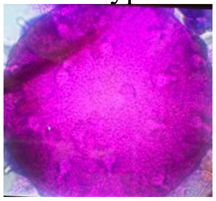 | Pollen Type 22<br>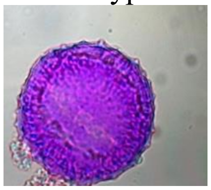 | Pollen Type 23<br>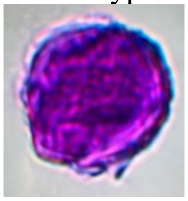 | Pollen Type 24<br>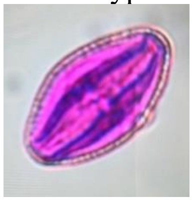 | Pollen Type 25<br>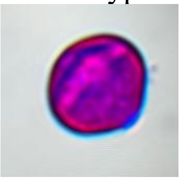 |

| | | | | |
|---|---|---|---|---|
| Pollen Type 26 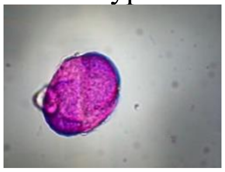 | Pollen Type 27 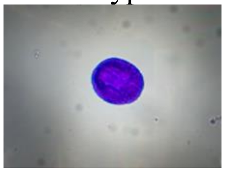 | Pollen Type 28 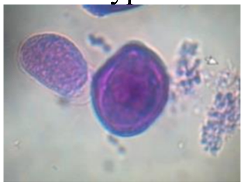 | Pollen Type 29 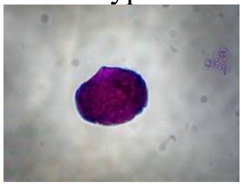 | Pollen Type 30 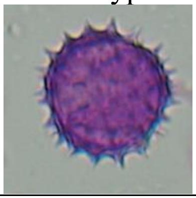 |
| Pollen Type 31 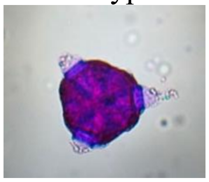 | Pollen Type 32 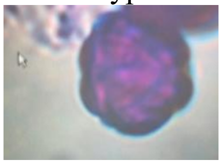 | Pollen Type 33 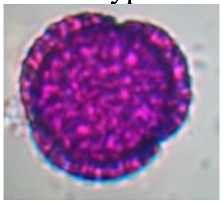 | Pollen Type 34 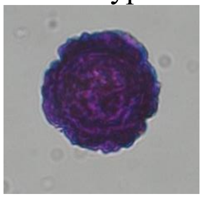 | Pollen Type 35 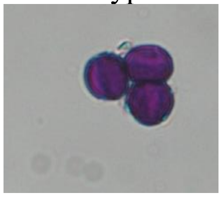 |
| Pollen Type 36 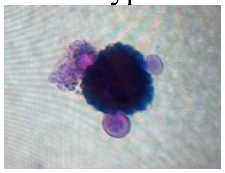 | Pollen Type 37 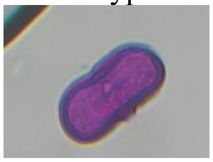 | Pollen Type 38 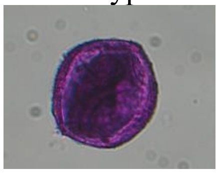 | Pollen Type 39 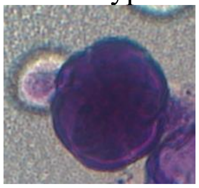 | Pollen Type 40 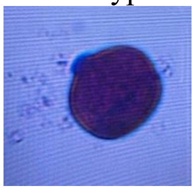 |

**Figure S2**: All of the bar plots from the pollen loads screened collected for individual bumble bee workers (*Bombus impatiens*) over ten days. The y-axis is the proportion of each pollen type, identified via microscopy, and the x-axis is the time of pollen collection in hours starting from time 0, which is the beginning of the respective trials. Each color represents a different pollen morphotype. For this figure, we used a 3% threshold as contamination (Louveaux et al. 1978); pollen designated as contamination was removed from the figures for visual clarity. Only bees with three or more pollen samples collected are represented from trial 1.

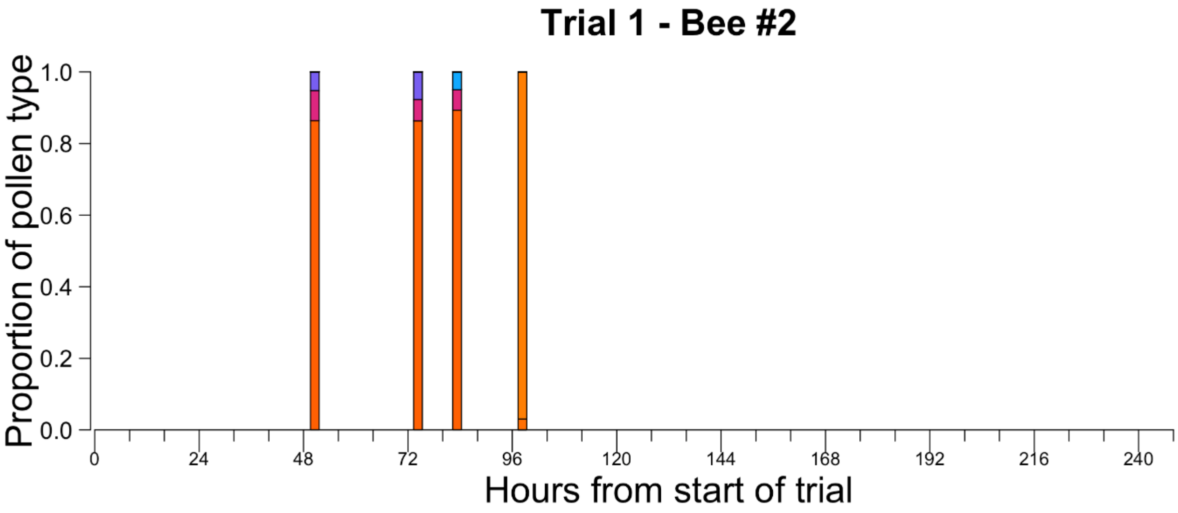

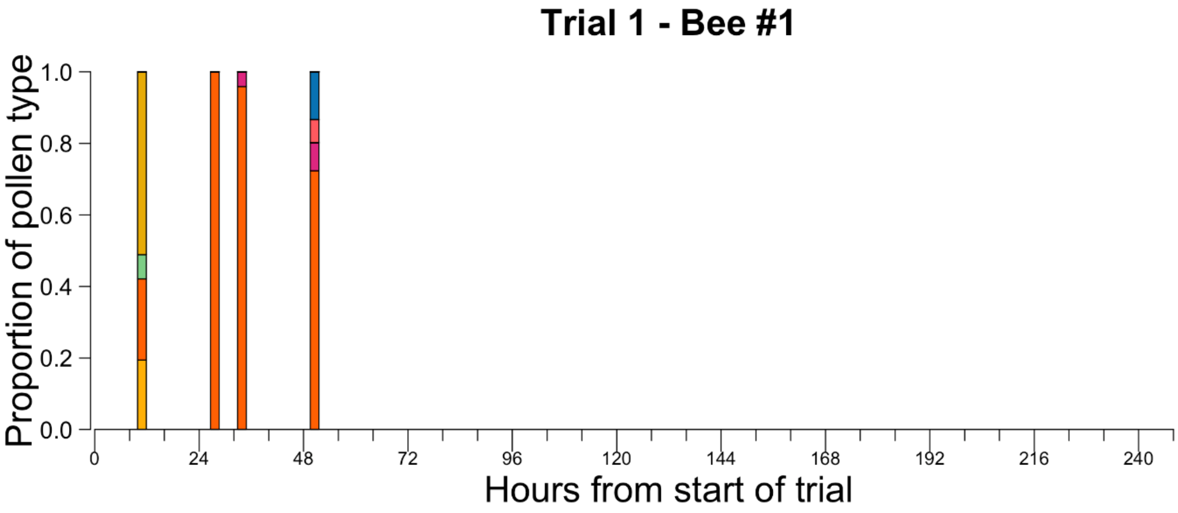

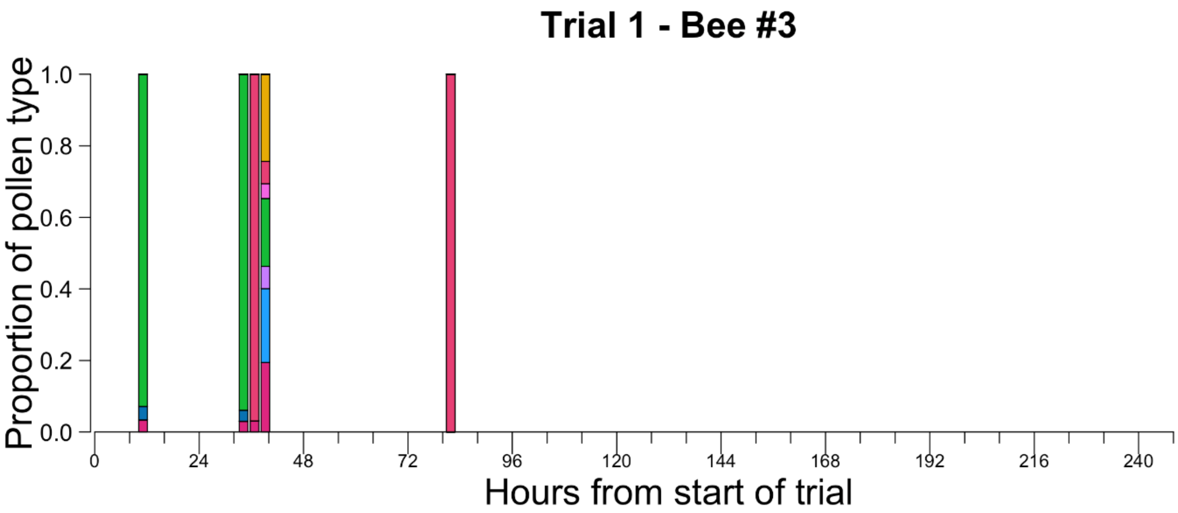
Trial 1 - Bee #3
Proportion of pollen type
1.0
0.8
0.6
0.4
0.2
0.0
0
24
48
72
96
120
144
168
192
216
240
Hours from start of trial

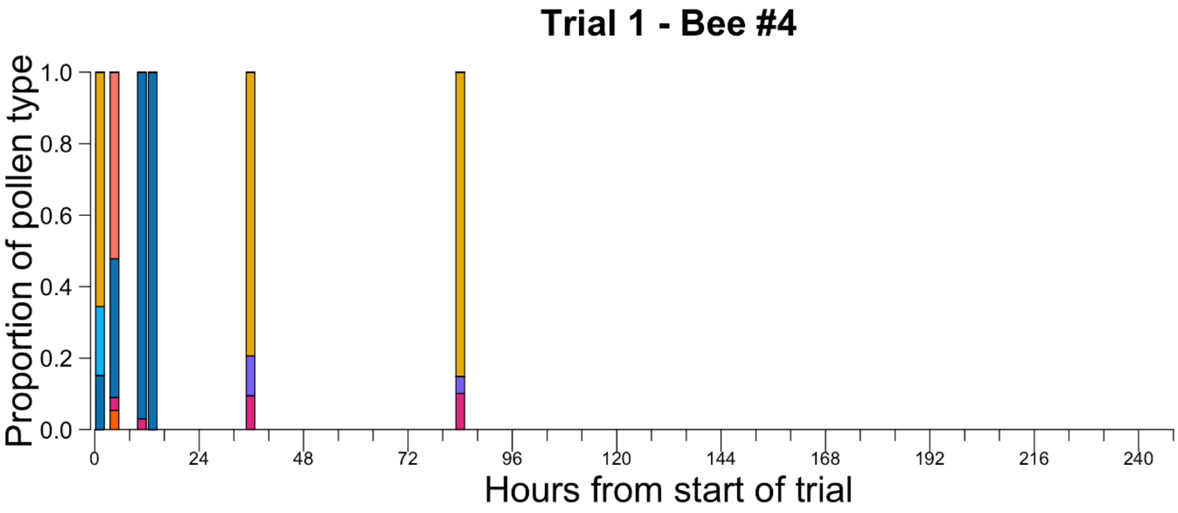
Trial 1 - Bee #4
Proportion of pollen type
1.0
0.8
0.6
0.4
0.2
0.0
0
24
48
72
96
120
144
168
192
216
240
Hours from start of trial

Trial 1 - Bee #5

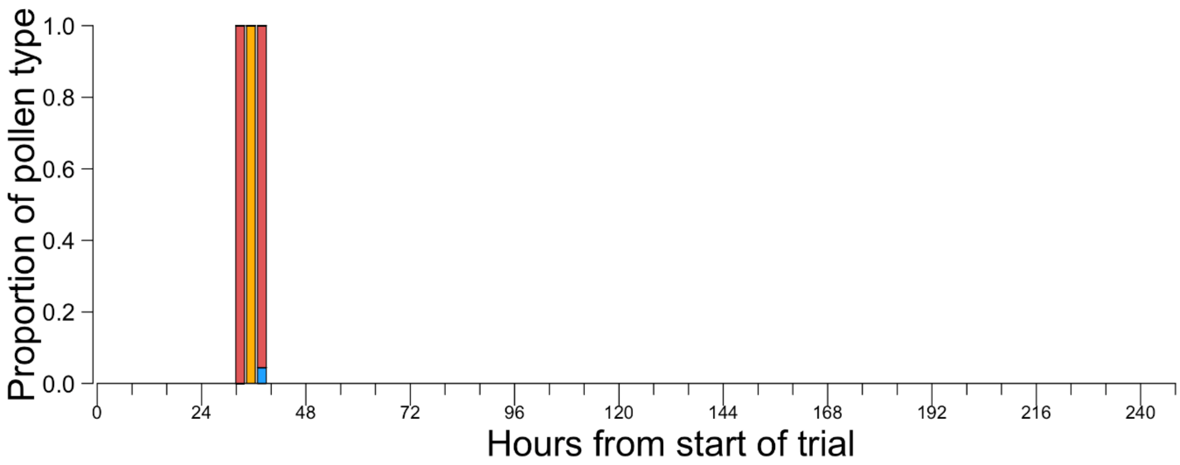

Trial 1 - Bee #6

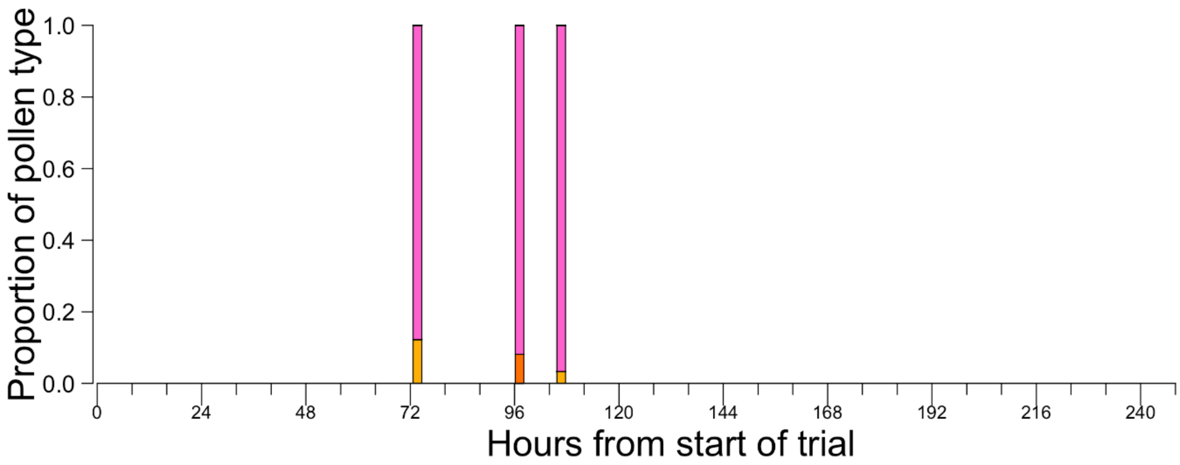

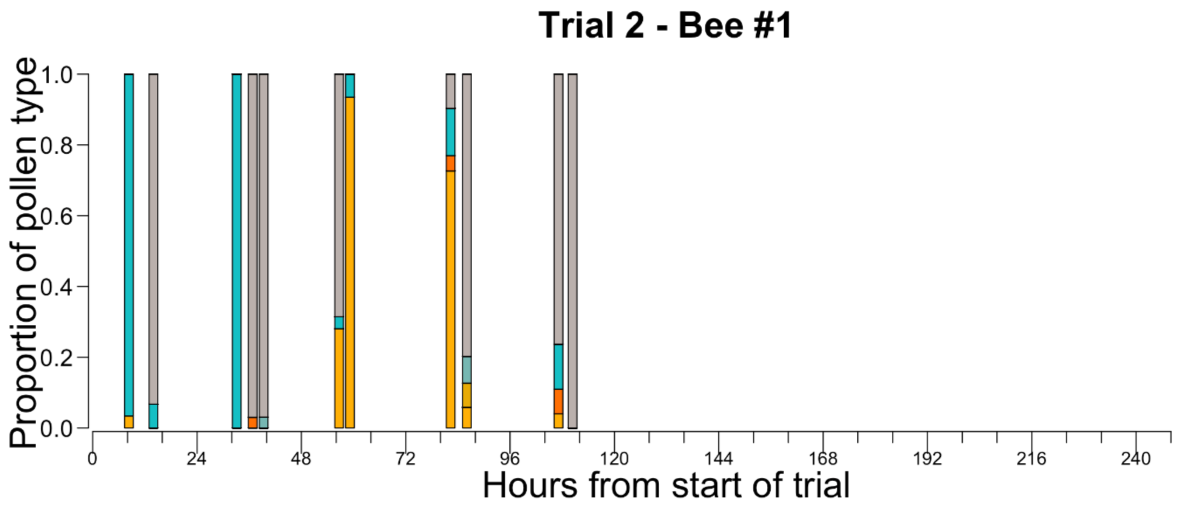
Trial 2 - Bee #1
Proportion of pollen type
1.0
0.8
0.6
0.4
0.2
0.0
0
24
48
72
96
120
144
168
192
216
240
Hours from start of trial

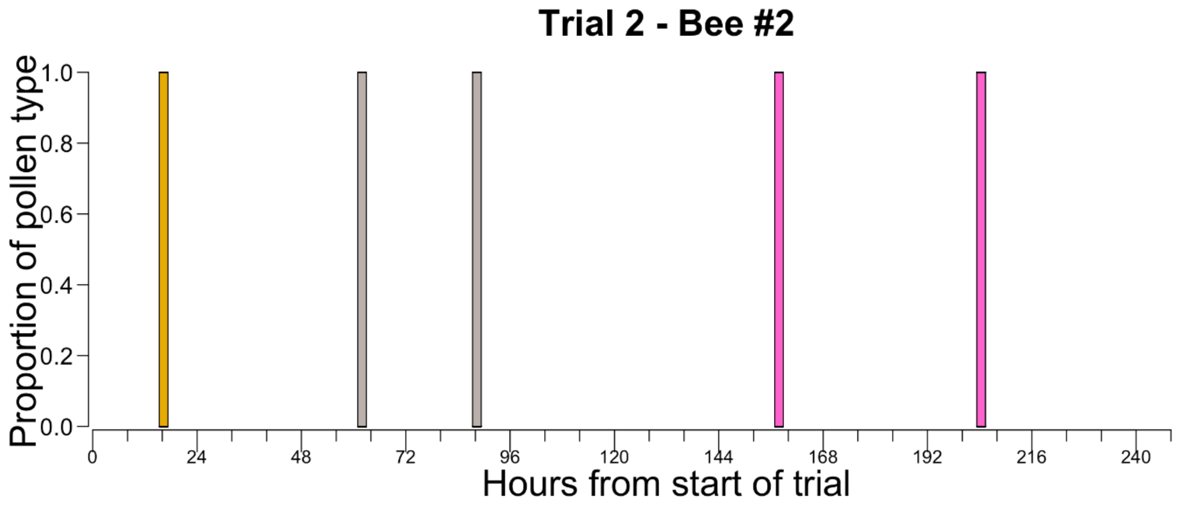
Trial 2 - Bee #2
Proportion of pollen type
1.0
0.8
0.6
0.4
0.2
0.0
0
24
48
72
96
120
144
168
192
216
240
Hours from start of trial

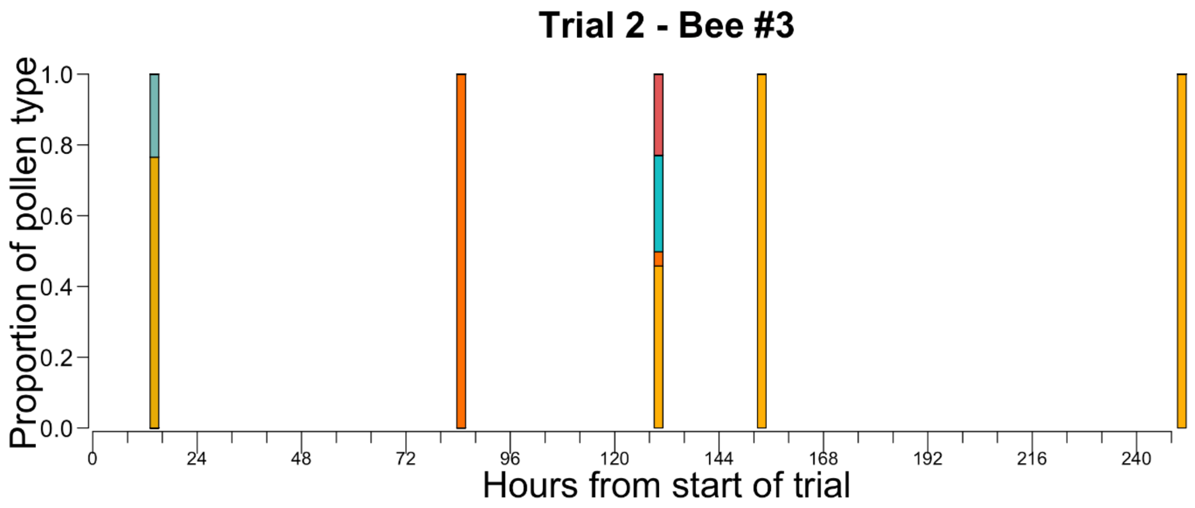
Trial 2 - Bee #3
Proportion of pollen type
0.0
0.2
0.4
0.6
0.8
1.0
0
24
48
72
96
120
144
168
192
216
240
Hours from start of trial

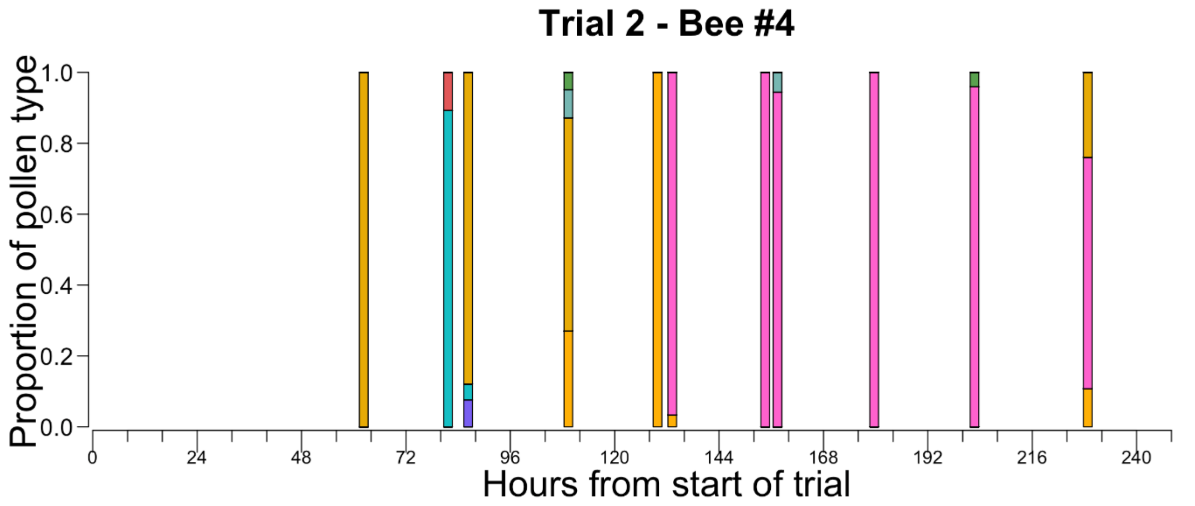
Trial 2 - Bee #4
Proportion of pollen type
0.0
0.2
0.4
0.6
0.8
1.0
0
24
48
72
96
120
144
168
192
216
240
Hours from start of trial

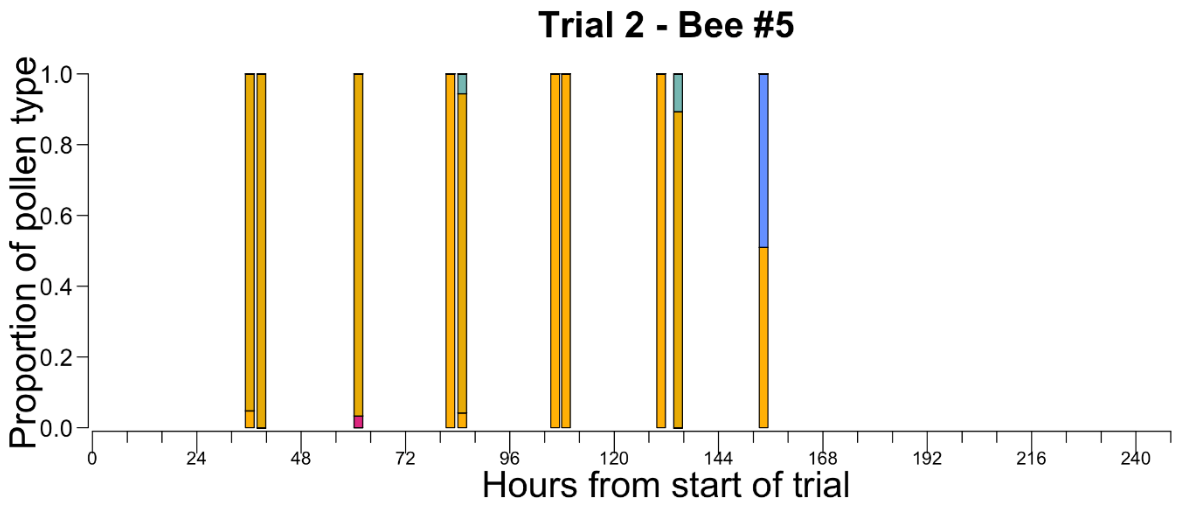

Trial 2 - Bee #5
Proportion of pollen type
0.0
0.2
0.4
0.6
0.8
1.0
0
24
48
72
96
120
144
168
192
216
240
Hours from start of trial

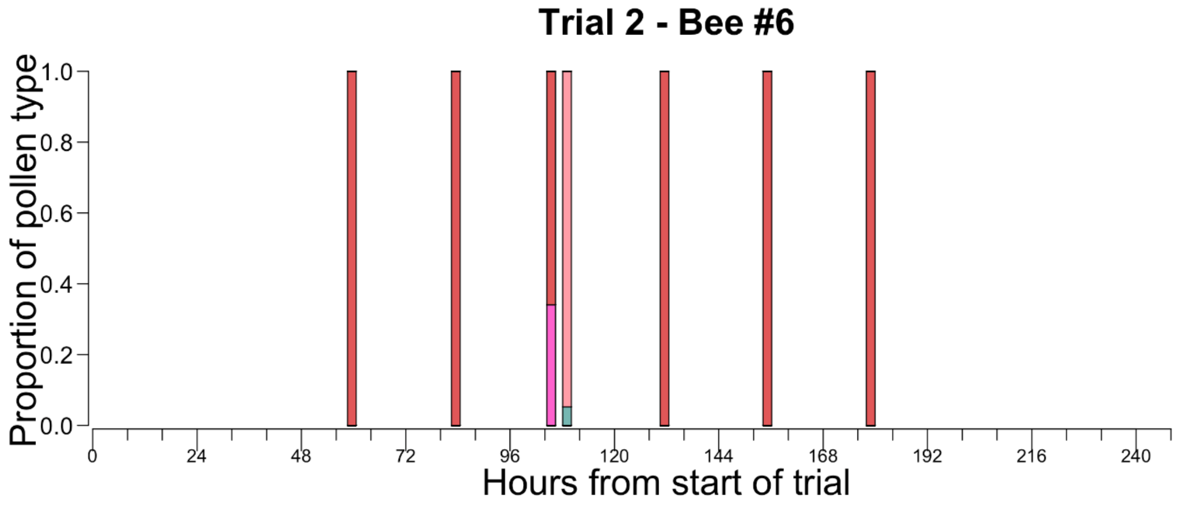

Trial 2 - Bee #6
Proportion of pollen type
0.0
0.2
0.4
0.6
0.8
1.0
0
24
48
72
96
120
144
168
192
216
240
Hours from start of trial

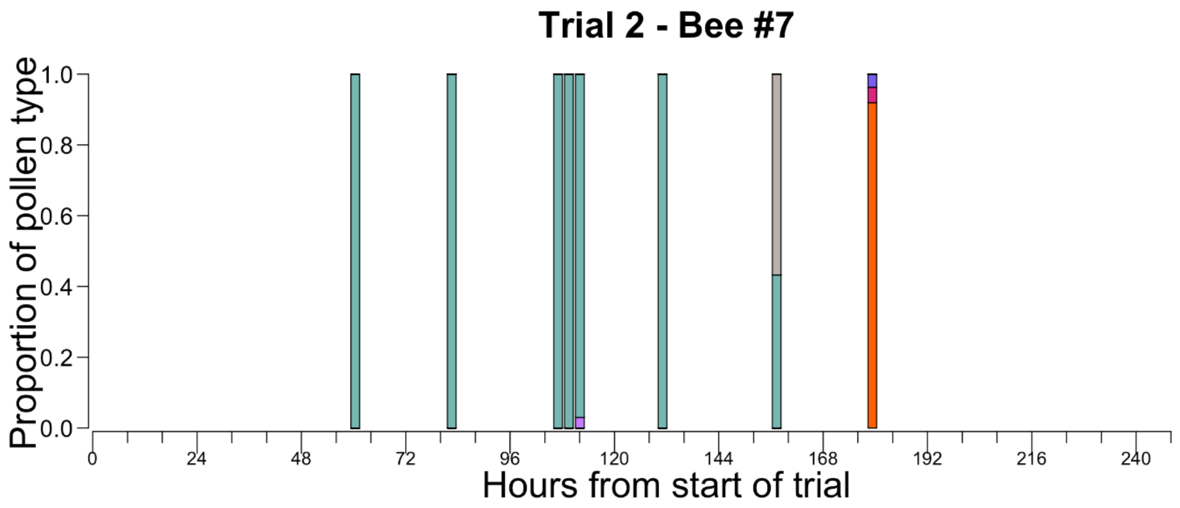
Trial 2 - Bee #7
Proportion of pollen type
0.0
0.2
0.4
0.6
0.8
1.0
0
24
48
72
96
120
144
168
192
216
240
Hours from start of trial

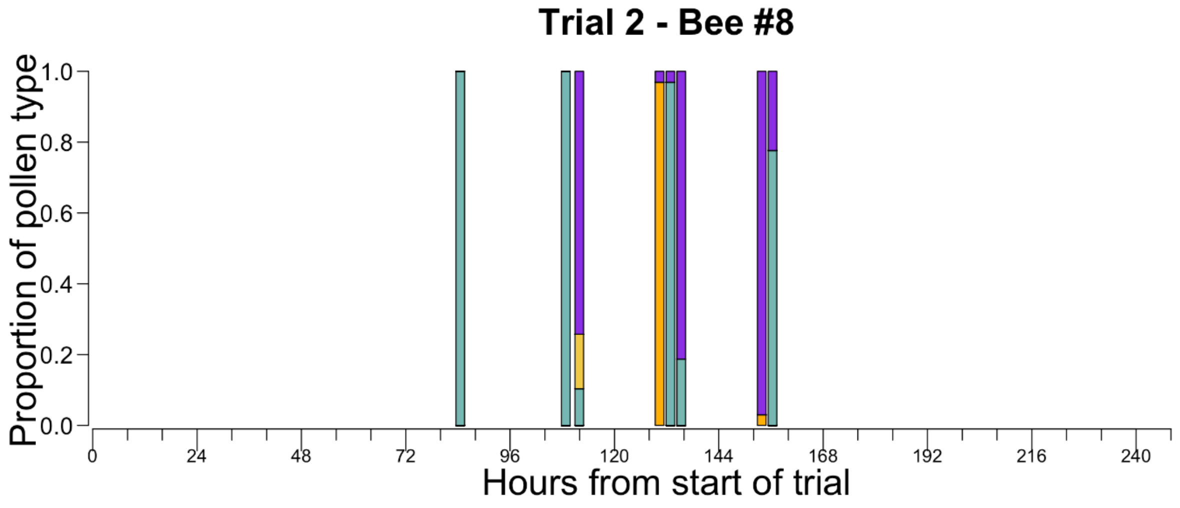
Trial 2 - Bee #8
Proportion of pollen type
0.0
0.2
0.4
0.6
0.8
1.0
0
24
48
72
96
120
144
168
192
216
240
Hours from start of trial

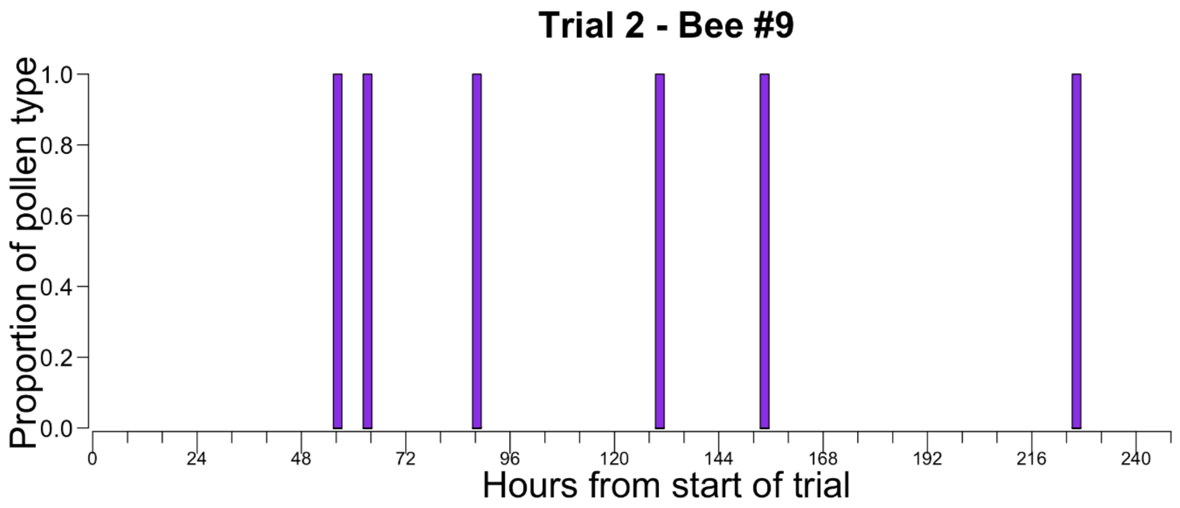
Trial 2 - Bee #9
Proportion of pollen type
1.0
0.8
0.6
0.4
0.2
0.0
0
24
48
72
96
120
144
168
192
216
240
Hours from start of trial

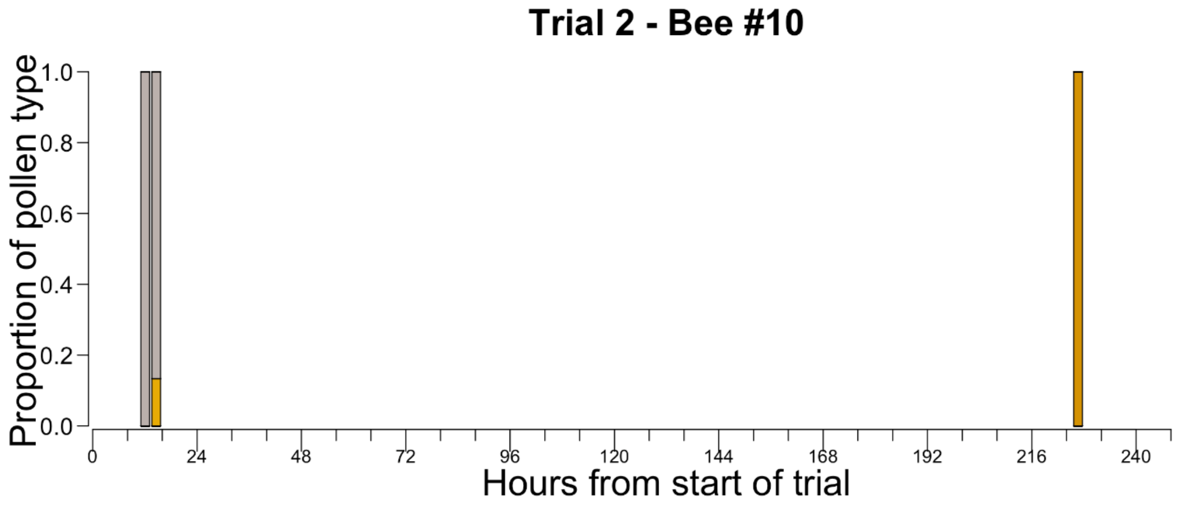
Trial 2 - Bee #10
Proportion of pollen type
1.0
0.8
0.6
0.4
0.2
0.0
0
24
48
72
96
120
144
168
192
216
240
Hours from start of trial

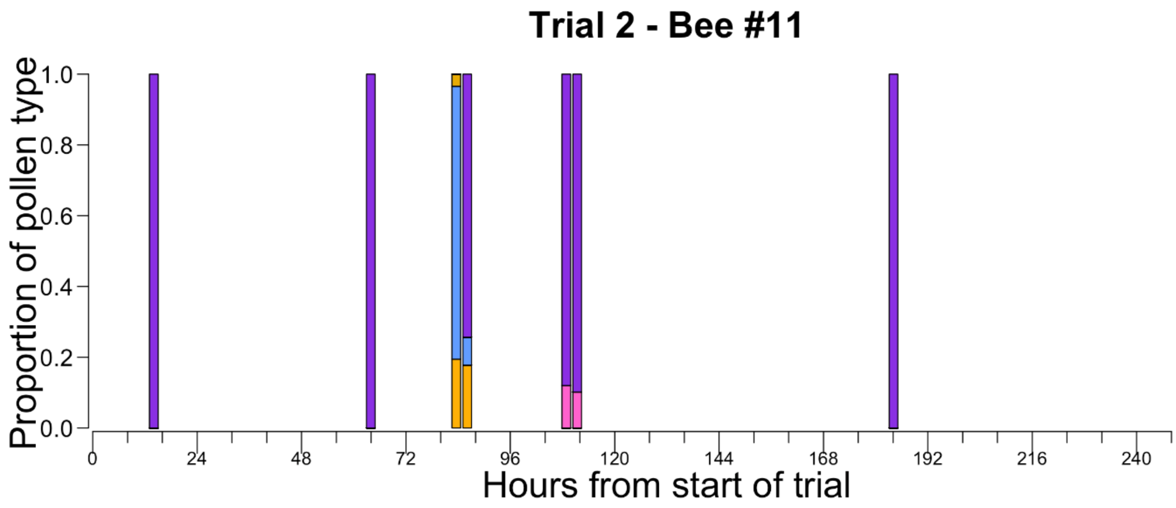
Trial 2 - Bee #11
Proportion of pollen type
1.0
0.8
0.6
0.4
0.2
0.0
0
24
48
72
96
120
144
168
192
216
240
Hours from start of trial

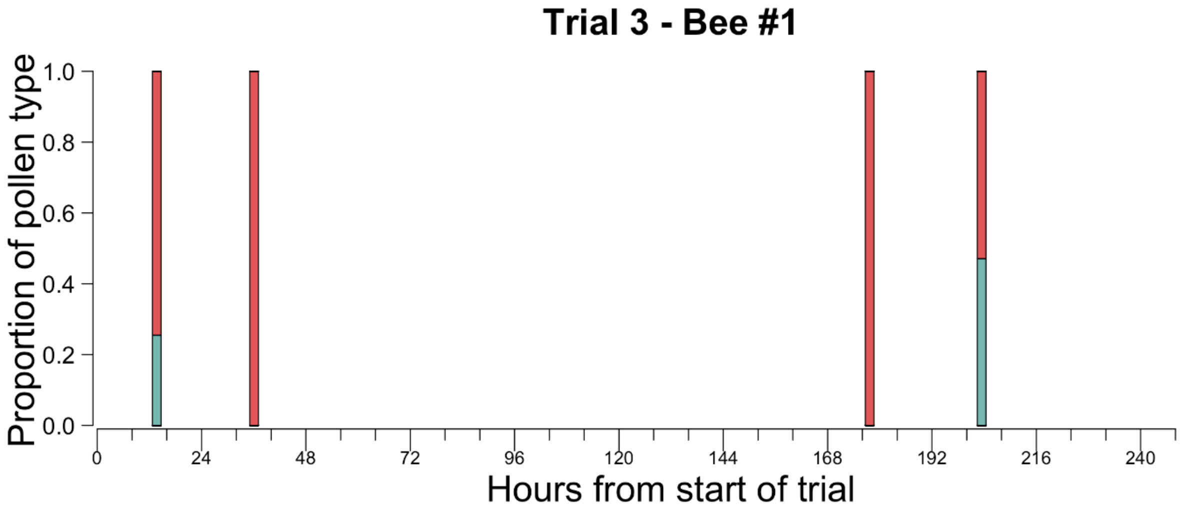
Trial 3 - Bee #1
Proportion of pollen type
1.0
0.8
0.6
0.4
0.2
0.0
0
24
48
72
96
120
144
168
192
216
240
Hours from start of trial

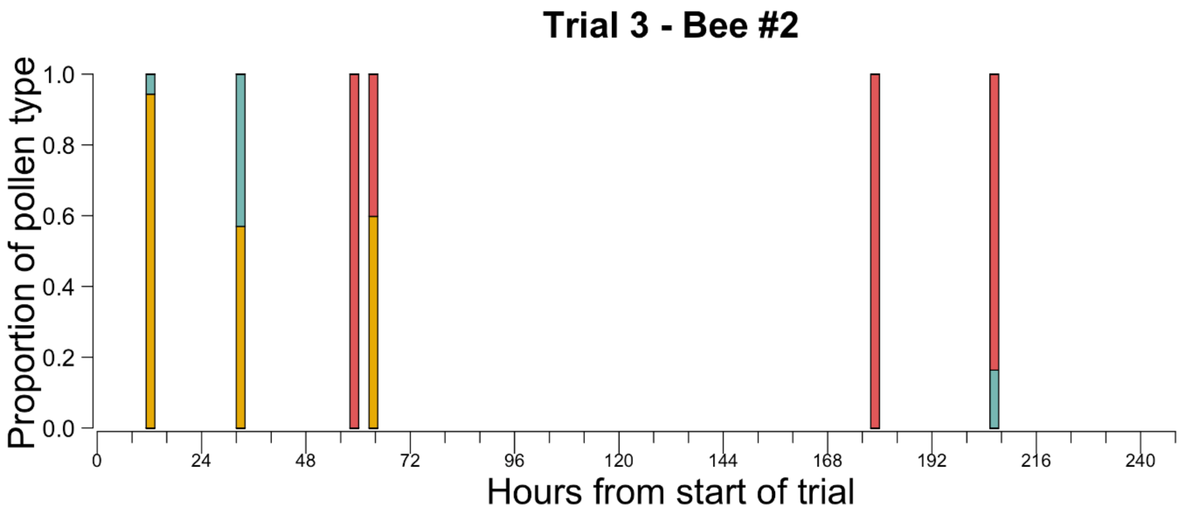

Trial 3 - Bee #2
Proportion of pollen type
1.0
0.8
0.6
0.4
0.2
0.0
0
24
48
72
96
120
144
168
192
216
240
Hours from start of trial

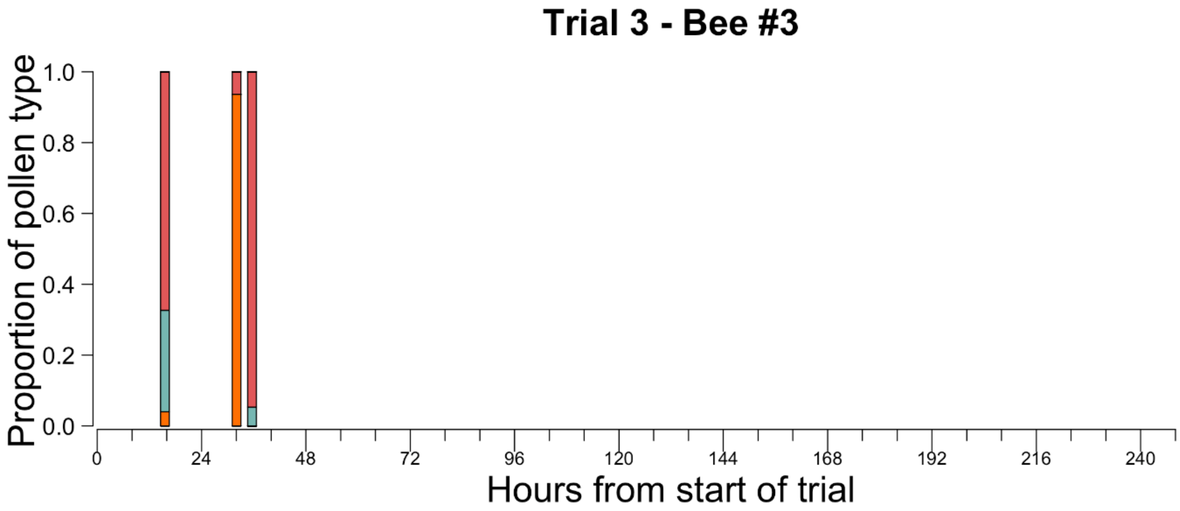

Trial 3 - Bee #3
Proportion of pollen type
1.0
0.8
0.6
0.4
0.2
0.0
0
24
48
72
96
120
144
168
192
216
240
Hours from start of trial

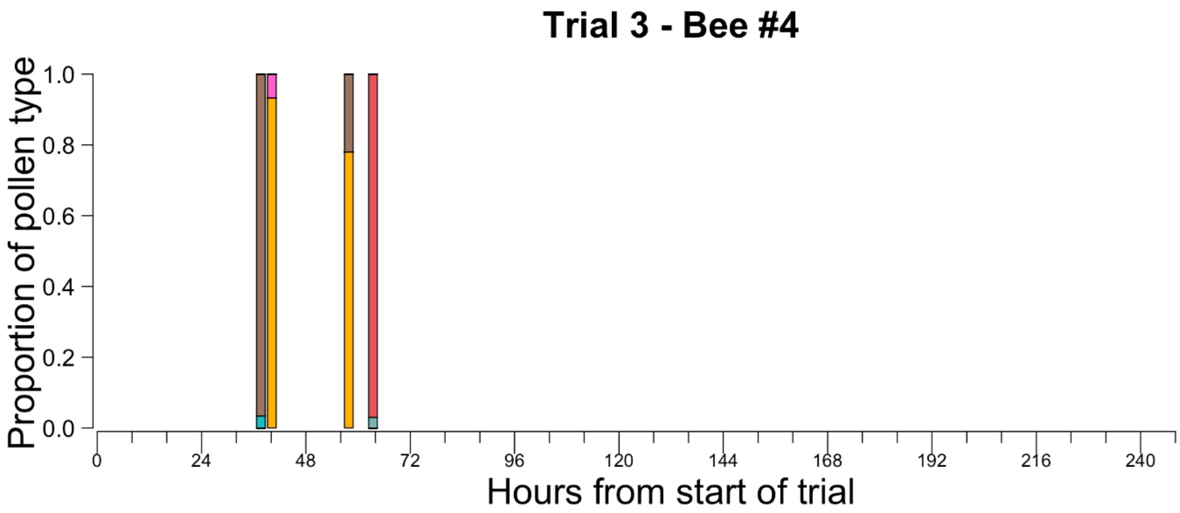
Trial 3 - Bee #4
Proportion of pollen type
0.0
0.2
0.4
0.6
0.8
1.0
0
24
48
72
96
120
144
168
192
216
240
Hours from start of trial

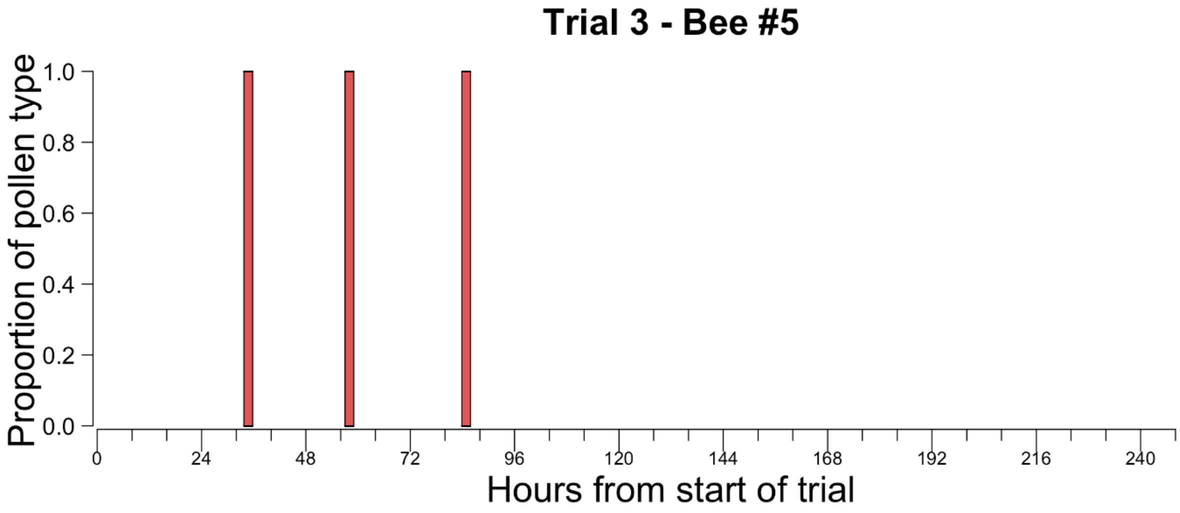
Trial 3 - Bee #5
Proportion of pollen type
0.0
0.2
0.4
0.6
0.8
1.0
0
24
48
72
96
120
144
168
192
216
240
Hours from start of trial

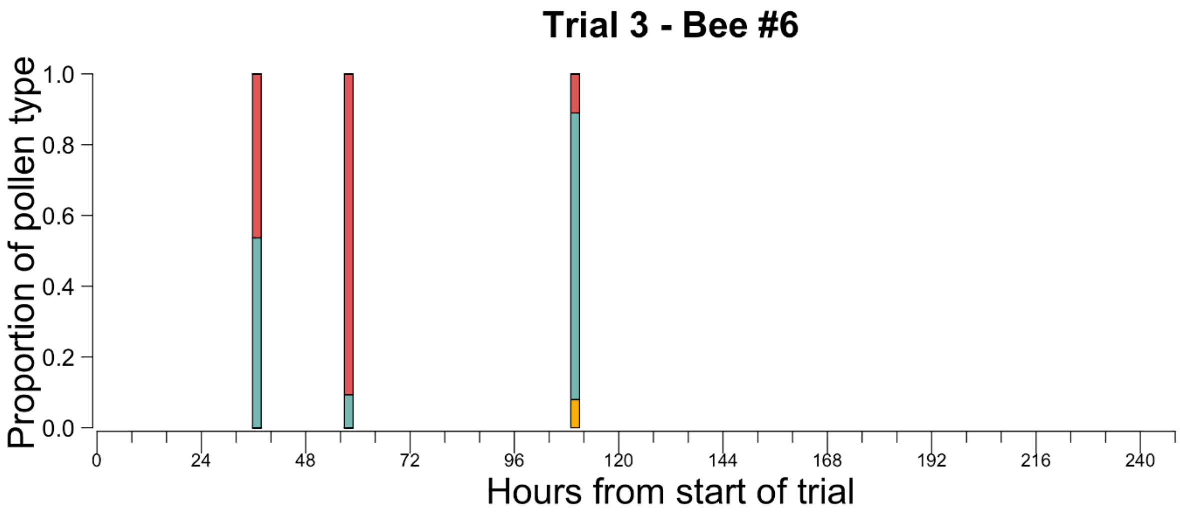
Trial 3 - Bee #6
Proportion of pollen type
1.0
0.8
0.6
0.4
0.2
0.0
0
24
48
72
96
120
144
168
192
216
240
Hours from start of trial

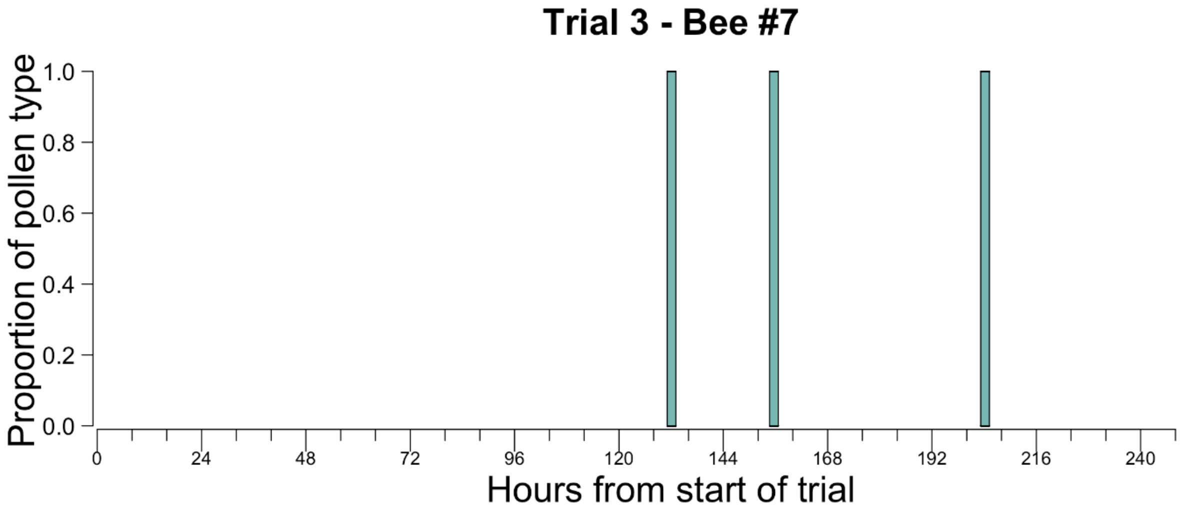
Trial 3 - Bee #7
Proportion of pollen type
1.0
0.8
0.6
0.4
0.2
0.0
0
24
48
72
96
120
144
168
192
216
240
Hours from start of trial

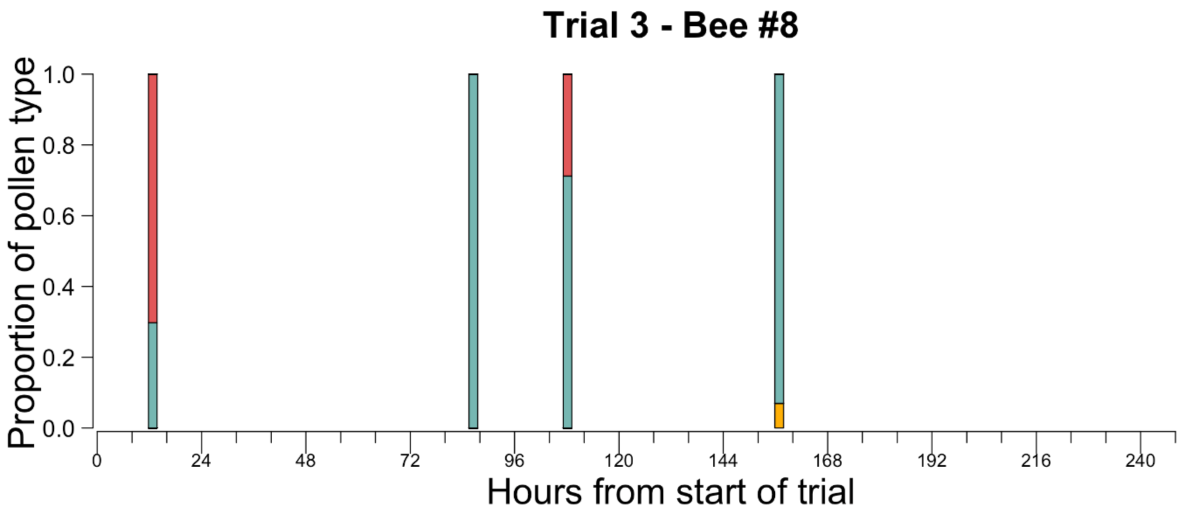

Trial 3 - Bee #8
Proportion of pollen type
1.0
0.8
0.6
0.4
0.2
0.0
0
24
48
72
96
120
144
168
192
216
240
Hours from start of trial

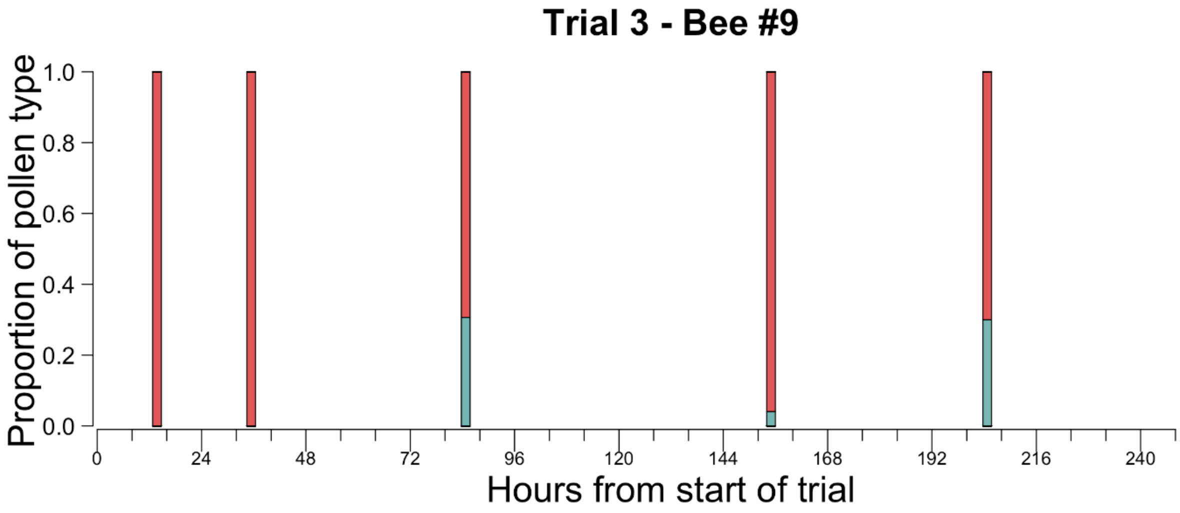

Trial 3 - Bee #9
Proportion of pollen type
1.0
0.8
0.6
0.4
0.2
0.0
0
24
48
72
96
120
144
168
192
216
240
Hours from start of trial

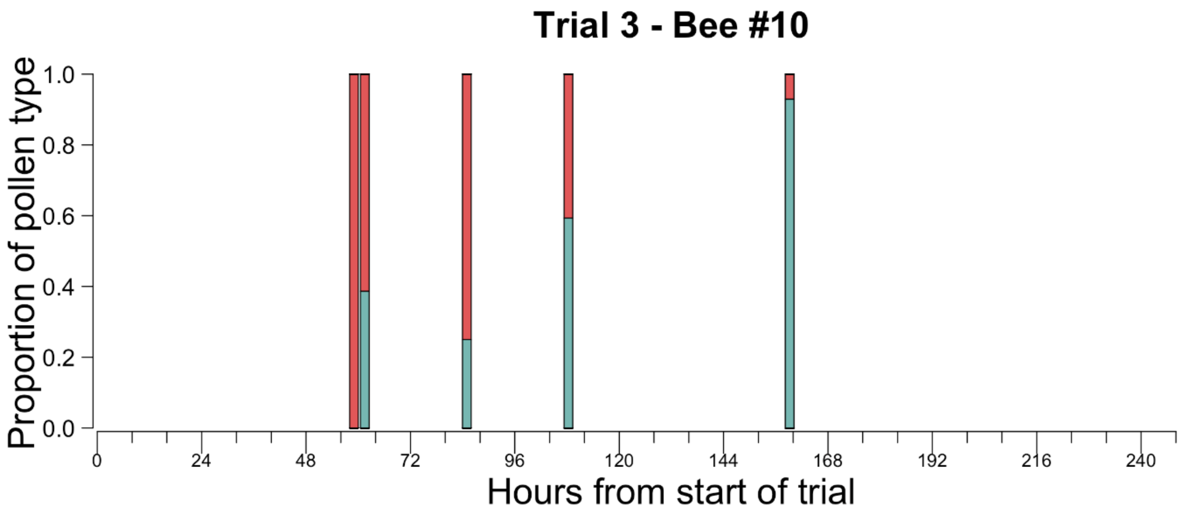
Trial 3 - Bee #10
Proportion of pollen type
1.0
0.8
0.6
0.4
0.2
0.0
0
24
48
72
96
120
144
168
192
216
240
Hours from start of trial

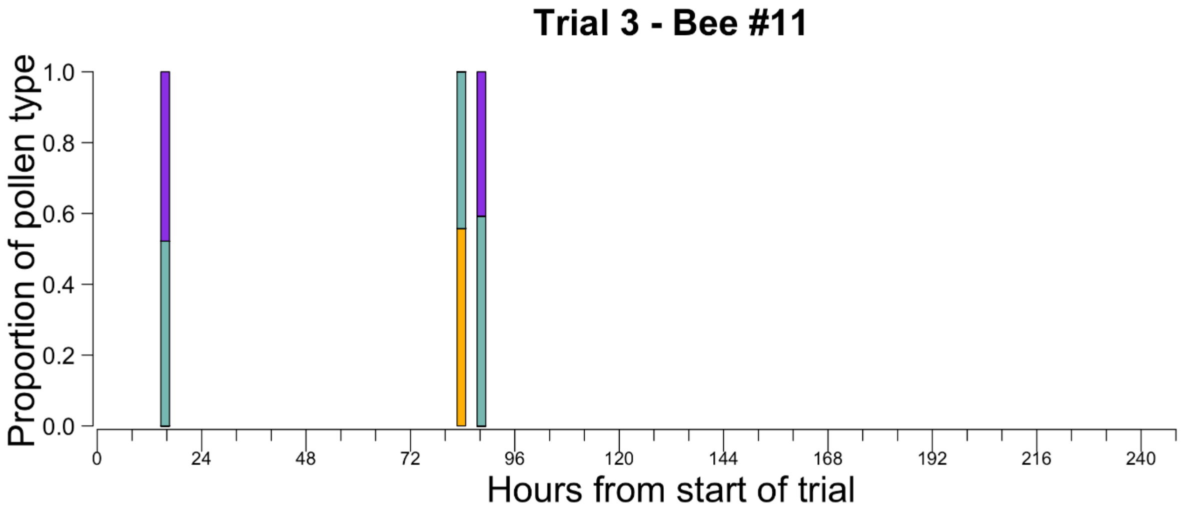
Trial 3 - Bee #11
Proportion of pollen type
1.0
0.8
0.6
0.4
0.2
0.0
0
24
48
72
96
120
144
168
192
216
240
Hours from start of trial

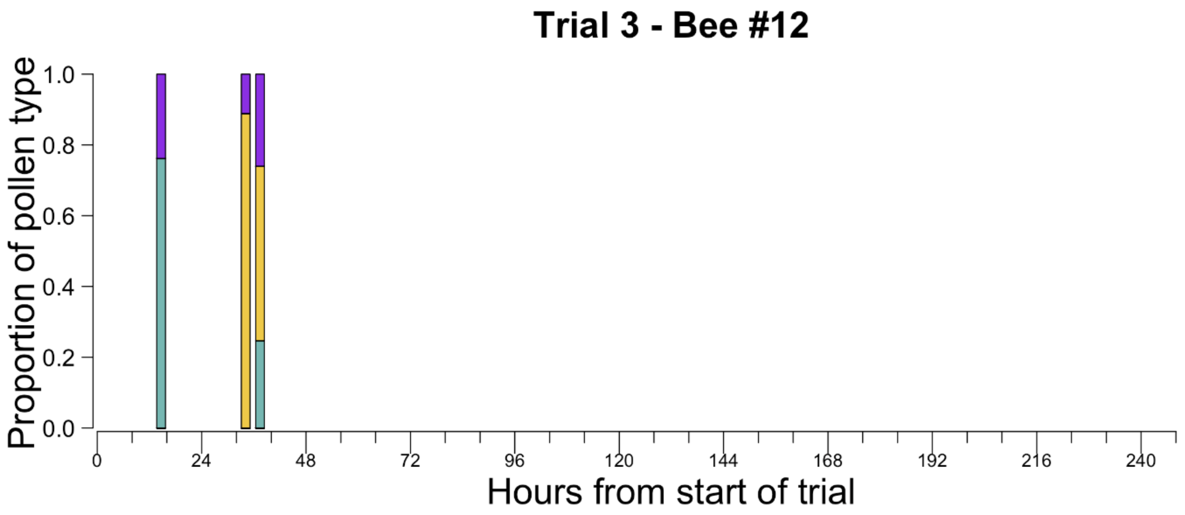

Trial 3 - Bee #12
Proportion of pollen type
1.0
0.8
0.6
0.4
0.2
0.0
0
24
48
72
96
120
144
168
192
216
240
Hours from start of trial

**Figure S3**: The histograms show the frequency of the most common pollen type and the second most common pollen type in each sample for each trial, shown separately.

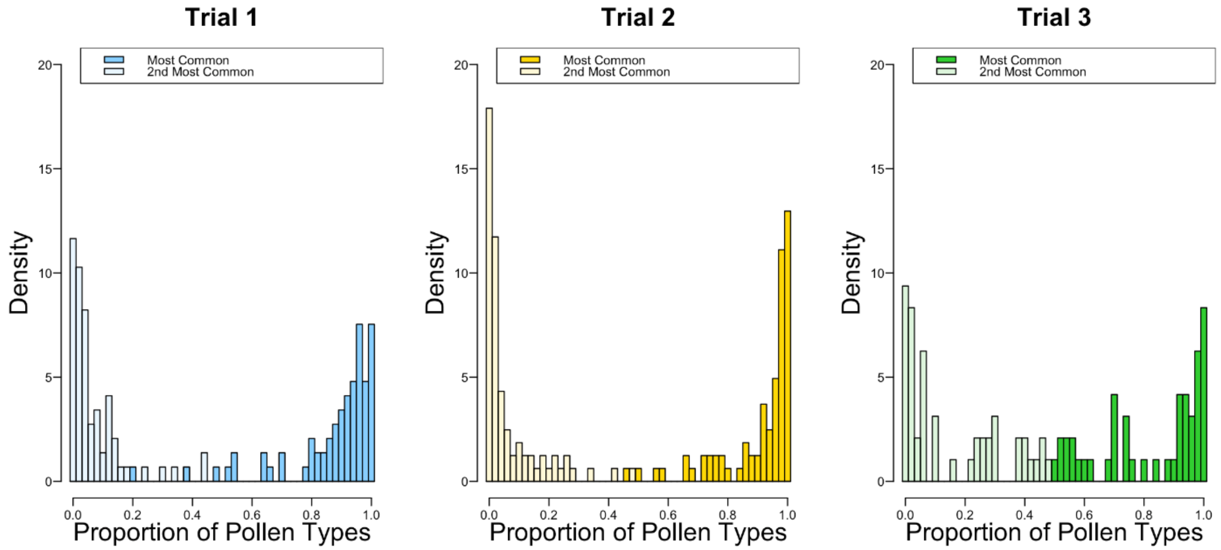

**Figure S4**: Simpson's diversity index as a measure of the effective number of pollen types. Each point corresponds to Simpson's index of all pollen samples from the same bee pooled within a moving window of 1, 2, 3, 4, and 5 days, respectively. The pink lines were calculated using randomly selected pollen samples from different bees and hence represent the equivalent Simpson's indices for the full colony's repertoire.

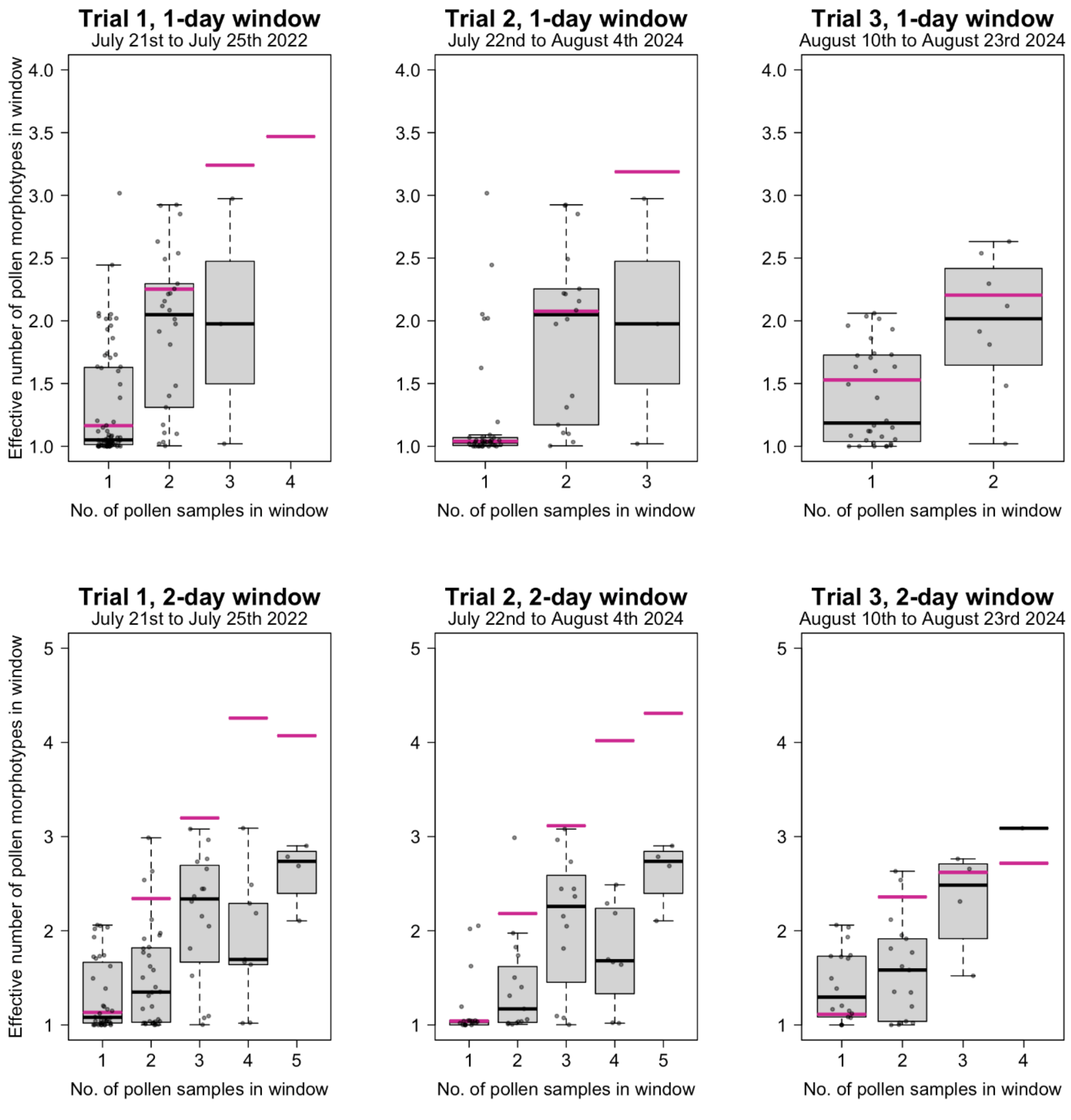

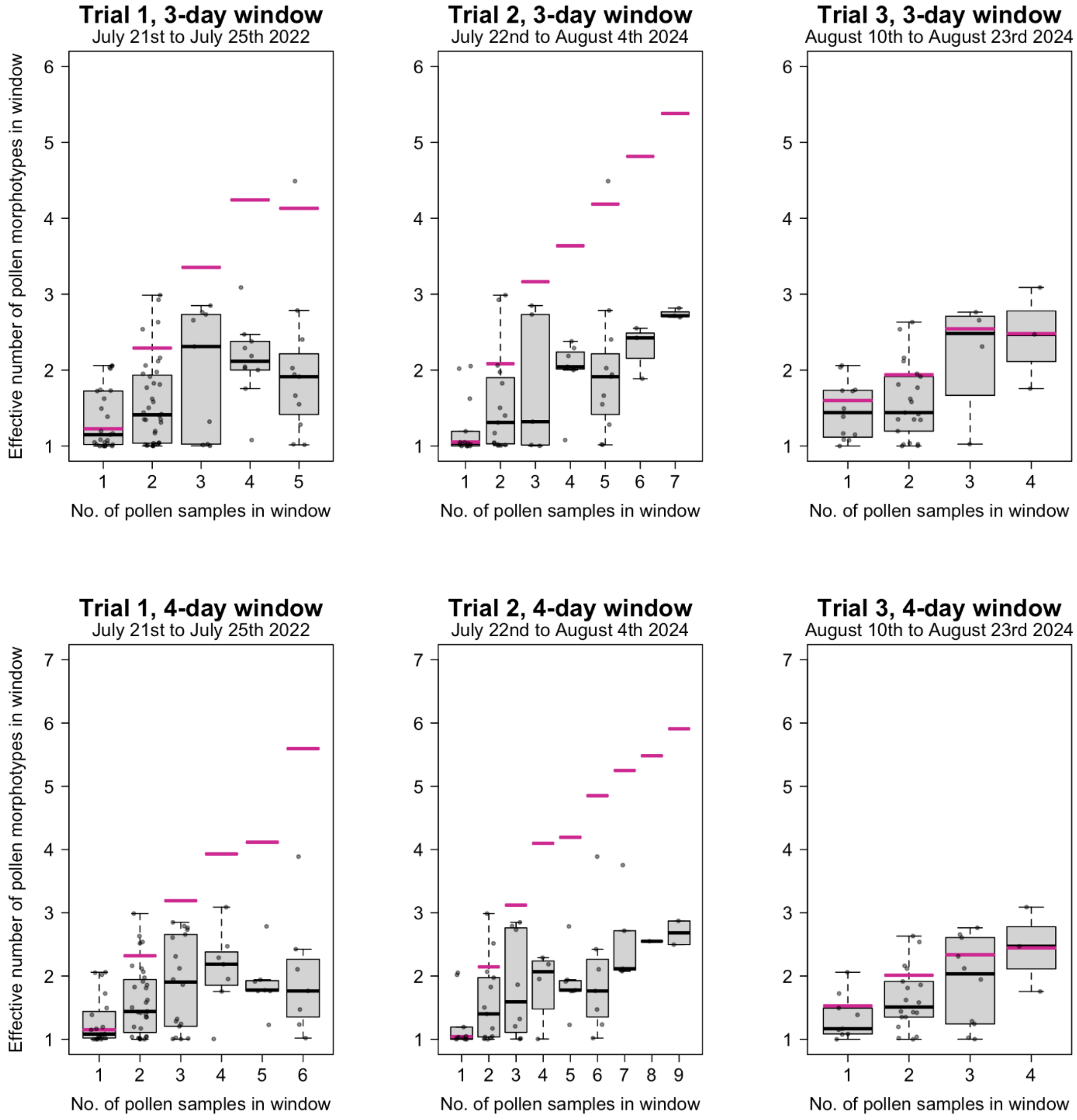

Trial 1, 3-day window
July 21st to July 25th 2022
Trial 2, 3-day window
July 22nd to August 4th 2024
Trial 3, 3-day window
August 10th to August 23rd 2024
Trial 1, 4-day window
July 21st to July 25th 2022
Trial 2, 4-day window
July 22nd to August 4th 2024
Trial 3, 4-day window
August 10th to August 23rd 2024
Effective number of pollen morphotypes in window
No. of pollen samples in window

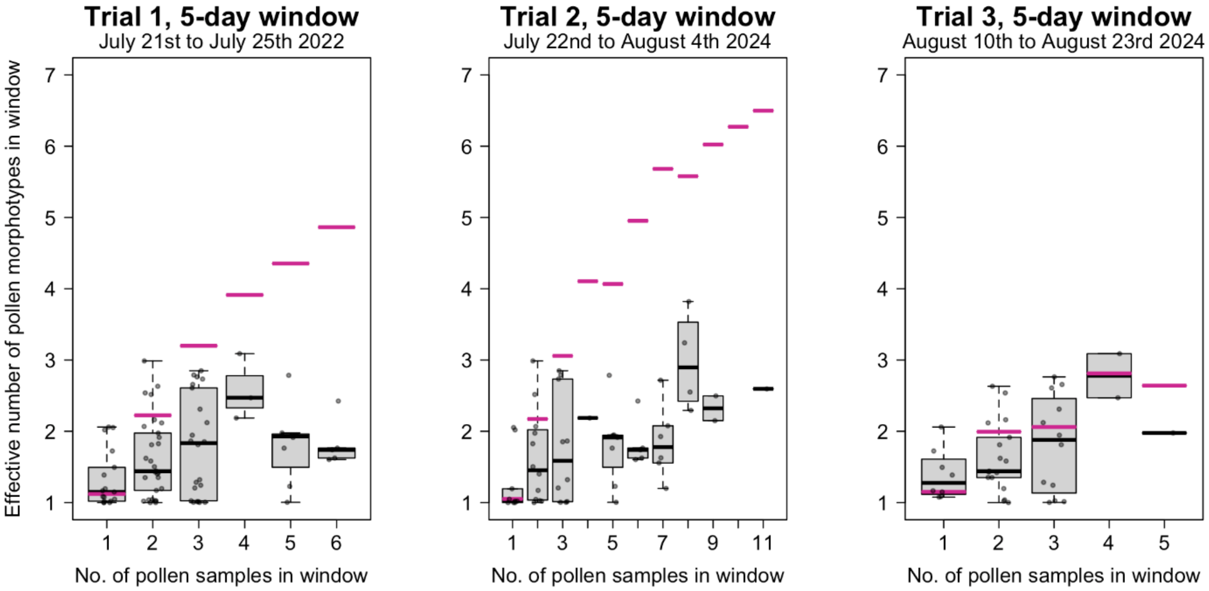

**Figure S5**: Drift did not have a significant effect on Jaccard similarity. Same as Figure 6 in the main text, except that for pollen sample pairs taken from the same bee in Trials 2 and 3, we differentiated between pairs where the bee switched colonies in between (see-green points) and those where the bee did not (grey points). This was to assess whether a bee that switched colony was more likely to change floral repertoire. For Trial 1, only one colony was used so switching between colonies was not possible. Because we did not find a significant difference between colony-switching vs. non-colony-switching pairs ($p > 0.05$), the best-fit line in black used both types of pairs.

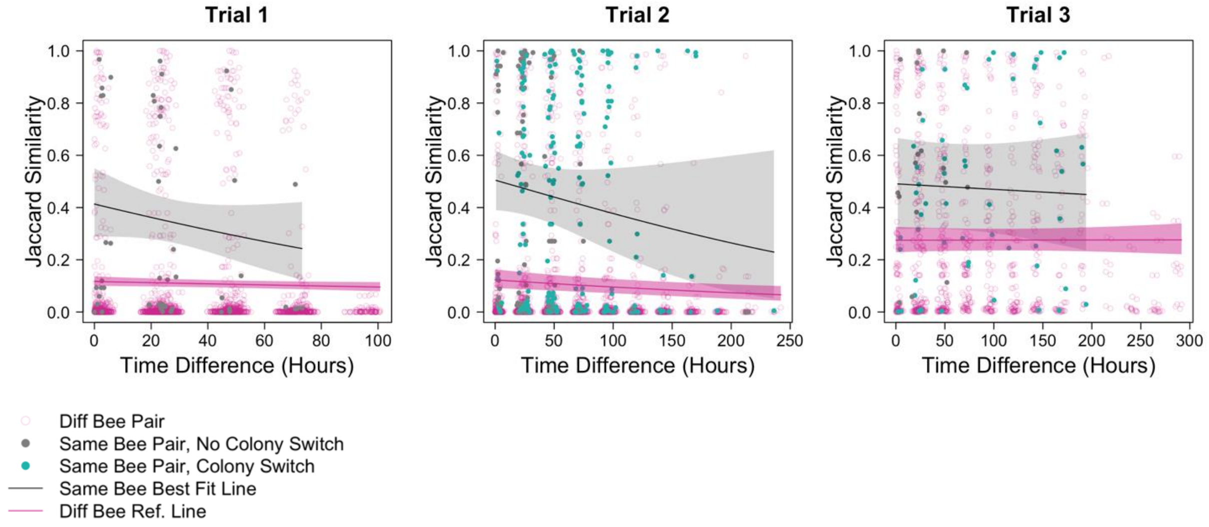

#### Section S1: Assessing the effects of colony drift on floral constancy

In our analysis involving the weighted Jaccard distance between pairs of pollen samples from the same bee or from different bees, a potential concern is the effects of drift during Trials 2 and 3, i.e., bees switching between the two colonies present. Could bees change repertoire as they switch between colonies? While drift is known to occur in nature, it was exacerbated in our experimental setup by the proximity of the two colonies in Trials 2 and 3, so any effects of drift on foraging repertoire would be amplified.

We began by testing whether same-bee pollen pairs collected from the same colony were more similar than same-bee pairs from different colonies. To do that we fitted an ordered beta GLMM with Jaccard similarity as the response, using data containing only same-bee pairs. The predictors were time separation and a "drift" factor variable with two levels indicating if the pollen pair came from the same or different colonies. We used bee identity as the random effects grouping variable, with a random effect structure that included a random intercept and slope for the effects of time separation. The effect of the factor variable was not statistically significantly different from zero (Trial 2: $z=-1.8$, $p_{1\text{-sided}} = 0.96$; Trial 3: $z=1.5$, $p_{1\text{-sided}} = 0.07$).

However, even for a same-bee pair collected from the same colony, it was possible for the bee to have switched colony and back in between, so we repeated this analysis, but this time using a stricter criterion for the "drift" factor variable where same-bee pairs were considered to have no drift only when all other pollen samples in between the pairs were also from the same colony. Again, we did not find a significant difference (Trial 2: $z=-1.6$, $p_{1\text{-sided}} = 0.94$; Trial 3: $z=0.77$, $p_{1\text{-sided}}=0.22$), so drift-related variables were excluded as covariates from all subsequent analyses.

#### Section S2: Permutation tests for Jaccard similarity analysis

When we used Jaccard similarities between pollen pairs as the response variables in GLMMs, the asymptotic distribution of the z-statistics from Wald tests may not be the normal distribution. This is because the similarities are not independent of one another: modifying one pollen sample simultaneously changes the similarities of all pollen pairs containing that sample. Therefore, for any significant results using the naive Wald test, we further confirmed the results using permutation tests to obtain sampling distributions of the z-score under the null hypotheses.

To test the effects of time separation between pollen pairs while still controlling for the same/different-bee binary predictor, we permuted the timings between samples with the same bee IDs. To test the effects of the binary predictor while still controlling for the time separation between pollen pairs, we permuted the bee IDs between samples.

We acknowledge that these permutation tests are not perfect. First, the exchangeability we have assumed in place of each null hypothesis is actually stricter than required by the hypothesis, because they also reduce some of the random effects variances to zero. Second, when testing the effects of the binary predictor, it is not possible to restrict the permutation of bee IDs to samples with the same timings, because timing is a continuous variable. Hence, while we still obtain permuted sets of data with the same likelihoods under the null hypothesis, they may differ in the predictor values. (For example, a certain time separation may be associated with two samples from the same bee in one permutation, and from two different bees in another permutation.) Therefore, the p-value from the permutation test is no longer a conditional p-value because we are allowing the predictor values to differ between permutations.

### Section S3: The effects of floral constancy on $R_0$ when each bee has a repertoire size of two

#### S3-1 Introduction

Floral constancy can affect the spread and persistence of bee pathogens that are transmitted indirectly via the shared use of flowers. Ellner et al. (2020) has shown that mathematical models that ignore floral constancy often severely underpredict the pathogen basic reproduction number $R_0$. This is because if floral constancy is lifelong, then bees that visit a flower species will never visit a different species, so they can only transmit to and be infected by other bees that visit the same species. This partitions the bipartite plant-pollinator transmission network into disconnected subnetworks each anchored by a flower species. Because some flower species necessarily have traits more conducive to pathogen transmission than the average, such a partitioning can increase the persistence of pathogens that would have otherwise died out, because infectious contacts in "conducive" subnetworks are not being wasted on the rest of the community.[1]

However, these results were based on a model which assumes that each bee is constant to only a single flower species at a time, i.e., a repertoire size of one. In contrast, our experimental results have shown that while *Bombus impatiens* workers exhibit floral constancy, the constancy is not to a single pollen type but to a repertoire of effective size between two and three (based on Simpson diversity). Hence, the goal of this supplement is to update the analysis in Ellner et al. (2020) to allow a repertoire size of two. Unlike the case of repertoire size one, because non-identical repertoires can now still exhibit some degree of overlap, the abovementioned partitioning of the plant-pollinator network is no longer as "clean" even when constancy is lifelong, so our goal is to assess the extent to which ignoring floral constancy still underpredicts $R_0$.

Because there is a continuum of possible ways by which a Simpson diversity of two can be achieved, we will consider two modified models that correspond to two extremes in this continuum. In the first model, the repertoire is heavily dominated by one flower species, but also contains all other species in the community in small but non-negligible proportions. We call this the **random sampling model**, because such a repertoire may arise if each bee has a single specialty but occasionally visits other species in the community at random (perhaps to help decide whether there is a more rewarding species to switch its specialty to), a behaviour

[1] Some readers may find this counterintuitive because earlier studies have shown that clustering can protect networks from disease spread (Eguíluz and Klemm, 2002). However, these studies are often based on network models where nodes correspond to individuals and where the number of contacts scales with the number of edges. In contrast, nodes in our models correspond to compartments (large well-mixed populations or sub-populations), and the total contact rate of a bee is treated as an empirically-determined parameter instead of something that scales with the number of edges. More edges simply spread the contacts among more flower species.

also reported in Heinrich (1976). In the second model, the repertoire contains only two species in equal proportions. Using the notion of majors and minors introduced in Heinrich (1976), we call this the **double-major model**. Both types of repertoires have been observed in our data.

#### S3-2 The single-major model (Ellner et al., 2020)

##### S3-2.1 Model description

For completeness, we begin with a short recap of the original model in Ellner et al. (2020), which we will refer to as the **single-major model**. In this model, bee pathogens are transmitted indirectly via flowers: an infected bee can contaminate a flower it has visited, which in turn can infect a subsequent visitor. While the original model involves $Q$ bee species and $K$ flower species, here for simplicity we will assume that $Q=1$, i.e., there is only a single bee species of interest as far as the pathogen dynamics is concerned. Despite being generalists at the species level, we assume that individual bees exhibit floral constancy with repertoire size one, so each bee specialises in only a single flower species. Because of this, instead of pooling all bees together, we use different state variables for bees with different specialties. More specifically, the state variables are $S_k$ and $I_k$ for the number of susceptible and infected bees that specialise in flower species $k$, and $U_k$ and $C_k$ for the number of uncontaminated and contaminated species-$k$ flowers, where $k \in \{1,2,...K\}$.

Pathogen transmision from a bee to species-$k$ flowers, and from a species-$k$ flowers to bees, occur at rates $\alpha_k$ and $\beta_k$ respectively, when a bee is foraging on species-$k$ flowers. Bees die at a per-capita rate $\mu$ if uninfected, and at an increased per-capita rate $\mu+\nu$ if infected. Infected bees can also recover at a per-capita rate $\gamma$, although they do not gain immunity against re-infection. Meanwhile, contaminated species-$k$ flowers decontaminate at a per-capita rate $\zeta_k$.

Bees switch specialty at a per-capita rate $\sigma$, and the probability of switching from species $j$ to $k \neq j$ is given by $a_{k,j}$ (so $\sum_{k \neq j} a_{k,j} = 1$). When the community is disease-free and at steady state, $M$ gives the total number of bees, and $\eta_k$ the fraction of bees that specialise in species $k$ (so $\sum_k \eta_k = 1$). Note that the specialty fractions $\{\eta_1, \eta_2, ...\eta_K\}$ are derived parameters that we will relate to other model parameters later on. We assume that new bees are recruited in the same fractions as well, so the birth rate of species-$k$ specialists is given by $M\mu\eta_k$. Finally, we assume that the total number of species-$k$ flowers remain constant at $N_k$.

The dynamic equations are

$$\begin{aligned}\frac{dS_k}{dt} &= M\mu\eta_k + \gamma I_k - \beta_k S_k \frac{C_k}{N_k} - (\mu+\sigma)S_k + \sigma\sum_{j\neq k} a_{k,j}S_j,\\ \frac{dI_k}{dt} &= \beta_k S_k \frac{C_k}{N_k} - (\mu+\nu+\gamma+\sigma)I_k + \sigma\sum_{j\neq k} a_{k,j}I_j,\\ \frac{dC_k}{dt} &= \alpha_k \frac{U_k}{N_k} I_k - \zeta_k C_k,\\ U_k &= N_k - C_k.\end{aligned} \tag{S1}$$

Because of our assumptions about newly recruited bees, $\{\eta_1,\eta_2,...\eta_K\}$ need to satisfy the implicit equation

$$\sum_{j\neq k} a_{k,j}\eta_j = \eta_k \tag{S2}$$

for all $k$ to become the disease-free steady-state specialty fractions. If we further assume that the switching probabilities are parametrised by a positive switching weight vector $\{w_1,w_2,...,w_K\}$ of unit sum such that

$$a_{k,j} = \frac{w_k}{1-w_j}, \tag{S3}$$

then it can be shown that

$$\eta_k = \frac{w_k(1-w_k)}{Z}, \quad Z = \sum_j w_j(1-w_j). \tag{S4}$$

To reduce the number of parameters, it is useful to rescale the state variables. Defining

$$s_k \equiv S_k/M, \quad y_k \equiv I_k/M, \quad c_k \equiv C_k/N_k, \quad \tilde{\alpha}_k = M\alpha_k/N_k, \tag{S5}$$

the rescaled dynamic equations are

$$\begin{aligned}\frac{ds_k}{dt} &= \mu\eta_k + \gamma y_k - \beta_k s_k c_k - (\mu+\sigma)s_k + \sigma\sum_{j\neq k} a_{k,j}s_j,\\ \frac{dy_k}{dt} &= \beta_k s_k c_k - (\mu+\nu+\gamma+\sigma)y_k + \sigma\sum_{j\neq k} a_{k,j}y_j,\\ \frac{dc_k}{dt} &= \tilde{\alpha}_k(1-c_k)y_k - \zeta_k c_k.\end{aligned} \tag{S6}$$

Finally, for convenience we define $\tau \equiv 1/(\mu+\nu+\gamma)$, which can be interpreted as the mean infectious lifetime of an infected bee. Table S3-1 summarises all the state variables and parameters in this model as well as the subsequent models.

##### S3-2.2 $R_0$ in the slow and fast switching limits

Let $R_0(\sigma)$ be the value of $R_0$ at a switching rate of $\sigma$. To demonstrate how ignoring floral constancy can cause us to underestimate the ability of a pathogen to spread and persist, we want to compare the values of $R_0(\sigma)$ between two limits: $\sigma \to 0$ (lifelong constancy) and $\sigma \to \infty$ (no constancy). Because $R_0(\sigma)$ is given by the spectral radius of the next-generation matrix (NGM), we need to derive the NGMs in both limits. While one can do so using more systematic approaches such as those described in Diekmann et al. (2010), we find it more instructive to derive them using intuitive arguments.

Because the plant-pollinator transmission network is bipartite, it is convenient to define one "generation" to be either flower-to-flower (via bees) or bee-to-bee (via flowers), and to derive the corresponding NGM. By a flower-to-flower NGM, we mean the $K \times K$ matrix whose $(j,k)$th element is the answer to the following question: if we start with one newly-contaminated species-$k$ flower, and monitor the bees infected by direct contact with that one flower before it decontaminates, and then the flowers contaminated by direct contact with those bees before they recover or die, how many species-$j$ flowers become contaminated? Likewise, we can also define a bee-to-bee NGM. Both NGMs will give the same values of $R_0(\sigma)$, because they describe the asymptotically exponential generation-to-generation growth rate of an initially infinitesimal infection.

###### S3-2.2.1 The slow switching limit $\sigma \to 0$

In the limit $\sigma \to 0$, bees are born with some specialty and are assumed to stick with it for life, with fraction $\eta_k$ having species $k$ as their specialty.

While the flower-to-flower and bee-to-bee NGMs are equally easy to derive in this limit, we will choose the former to facilitate comparison with the double-major model later on. We start the rescaled model Eqn. (S6) with a very low level of infection in flower species $j$ (so $c_k(0) = \epsilon$ for $k = j$ and 0 otherwise) and all bees susceptible (so $s_k(0) = \eta_k$ for all $k$). The initially-contaminated species-$j$ flowers remain contaminated for an average duration of $1/\zeta_j$, producing new infections $y_j = \epsilon\beta_j\eta_j/\zeta_j$ among bees with specialty $j$. Each of those bees remains infected and alive for time $\tau$ on average, giving rise to $\tau\tilde{\alpha}_j$ contaminated flowers in species $j$. Because these bees never change speciality, they cannot give rise to contamination in species

Table S3-1: State variables, parameters/parameter combinations, and their definitions. Model "s" corresponds to the Ellner et al. (2020) single-major model, "r" the random sampling model, and "d" the double-major model.

| Parameter | Definition or formula | Units | Model(s) |
|---|---|---|---|
| $M$ | Total population of bees at the disease-free equilibrium | individuals | s, r, d |
| $b$ | Total birth rate of bees | bees/day | s, r, d |
| $S_k$ / $S_{\{j,k\}}$ | Total number of susceptible bees with specialty $k$ / repertoire $\{j,k\}$. | individuals | s, r / d |
| $I_k$ / $I_{\{j,k\}}$ | Total number of infected bees with specialty $k$ / repertoire $\{j,k\}$ | individuals | s, r / d |
| $N_k$ | Total population of species $k$ flowers | floral units | s, r, d |
| $C_k$ | Total number of contaminated species $k$ flowers | floral units | s, r, d |
| $\alpha_k$ | Transmission rate from an infected bee to non-contaminated flowers of species $k$, when the bee is foraging on flower species $k$. | (floral units) /bee /day | s, r, d |
| $\beta_k$ | Transmission rate from contaminated flower of species $k$ to susceptible bee with flower species $k$, when the bee is foraging on flower species $k$. | bees /(floral unit) /day | s, r, d |
| $\gamma$ | Bee rate of recovery from infection | day$^{-1}$ | s, r, d |
| $\mu$ | Death rate of susceptible bees. | day$^{-1}$ | s, r, d |
| $\nu$ | Additional death rate of infected bees. | day$^{-1}$ | s, r, d |
| $\tau$ | $1/(\gamma+\mu+\nu)$, mean time to bee death or recovery | days | s, r, d |
| $\zeta_k$ | Flower rate of recovery from infection. | day$^{-1}$ | s, r, d |
| $\sigma$ | Overall rate of switching. | day$^{-1}$ | s, r, d |
| $a_{m,j}$ / $a_{m\|\{j,k\}}$ | Switching probability to flower species $m$ for a bee with current specialty $j$, $m \neq j$ / current repertoire $\{j,k\}$, $m \notin \{j,k\}$ | unitless | s, r / d |
| $w_m$ | Switching weights, strictly positive with $\sum_{m=1}^{K} w_m = 1$ | unitless | s, r, d |
| $v_m$ | Sampling weights, strictly positive with $\sum_{m=1}^{K} v_m = 1$ | unitless | r |
| $\phi_k$ | Fraction of foraging trips that a bee with specialty $k$ performs random sampling of the floral community | unitless | r |
| $\rho_k$ | Fraction of bees with species $k$ in its repertoire | unitless | d |
| $\eta_k$ / $\eta_{\{j,k\}}$ | Equilibrium fraction of specialty $k$ / repertoire $\{j,k\}$ in the absense of disease | unitless | s, r / d |

Note: units correspond to Table 1 in Truitt et al. (2019), which gives estimates and ranges for many of the parameters for old field communities in upstate New York, USA.

$k \neq j$. Using $\mathbf{Z}(\sigma)$ to denote the flower-to-flower NGM with switching rate $\sigma$, we therefore have

$$\mathbf{Z}(0)_{k,j} = \begin{cases} \tau \tilde{\alpha}_j \beta_j \eta_j / \zeta_j, & k = k, \\ 0, & \text{otherwise.} \end{cases} \tag{S7}$$

Note that the NGM is diagonal: contamination in species-$j$ flowers can only spread to other species-$j$ flowers. Hence, as stated in the introduction, the plant-pollinator network has been partitioned into disconnected subnetworks each anchored by a single flower species. Each diagonal element can be interpreted as $R_0(0)$ of the corresponding subnetwork. Because the pathogen can persist in the community as long as it can do so in at least one subnetwork, this implies that the community $R_0(0)$ is given by the largest of the subnetwork $R_0(0)$, so

$$R_{0,s}(0) = \tau \max_k (\tilde{\alpha}_k \beta_k \eta_k / \zeta_k) \tag{S8}$$

the subscript $s$ indicating that formula is for the single-major model.

###### S3-2.2.2 The fast switching limit $\sigma \to \infty$

In the limit $\sigma \to \infty$, there is really only one type of bee, a generalist that visits every flower species in accordance to the fractions $\{\eta_1, \eta_2, ... \eta_K\}$. Therefore, the bee-to-bee NGM has dimension one and is in fact equal to $R_0(\infty)$.

We can write down $R_0(\infty)$ explicitly using the following arguments. Once infected, a bee remains infectious for an average duration of $\tau$ until it recovers or dies, and therefore spends an average duration of $\tau \eta_j$ contaminating species-$j$ flowers. This produces $\tau \eta_j \tilde{\alpha}_j$ contaminated species-$j$ flowers (in the units of the rescaled model). Each such contaminated flower remains contaminated for average duration of $1/\zeta_j$, resulting in a time-integrated total force of infection of $\tau \eta_j \tilde{\alpha}_j \beta_j / \zeta_j$ from all first-generation contaminated species-$j$ flowers combined. Uninfected bees are exposed to this force of infection only while they are visiting species $j$, so the resulting number of second-generation bee infections is $\tau \eta_j^2 \tilde{\alpha}_j \beta_j / \zeta_j$. Summing across all flower species gives the overall bee-to-flower-to-bee transmission

$$R_{0,s}(\infty) = \tau \sum_{j=1}^{K} \eta_j^2 \tilde{\alpha}_j \beta_j / \zeta_j. \tag{S9}$$

#### S3-3 The random sampling model

The random sampling model differs from the Ellner et al. (2020) single-major model in that even though each bee still has a single specialty, it now goes on occasional random sampling trips, hence increasing its effective repertoire size.

For a species-$k$ specialist, we define $\phi_k$ as the fraction of foraging trips that are random sampling trips, during which the bee will visit every species except $k$ with relative frequency based on a sampling weight vector $\{v_1,v_2,...,v_K\}$ of unit sum. This means that a species-$k$ specialist bee will spend a fraction $(1-\phi_k)$ of its time visiting species $k$, and a fraction $\phi_k v_j/(1-v_k)$ of its time visiting species $j$ for each $j \neq k$. To achieve an effective repertoire size (Simpson diversity) of two, we require that $\phi_k$ satisfy

$$(1-\phi_k)^2+\phi_k^2\sum_{j\neq k}\left(\frac{v_j}{1-v_k}\right)^2=\frac{1}{\text{Simpson diversity}}=\frac{1}{2}. \tag{S10}$$

As an illustration, we simulated random weight vectors by drawing each component from a uniform distribution and then normalising the vector to have a unit sum. For $K=8$, we find that $\phi_k \simeq 0.3$ for all $k$ (there is surprisingly very little variation among $k$ even though the smallest and largest weights can differ by orders of magnitude in the simulations), so each bee will spend 70% of its time visiting its specialty, and 4.3% of its time visiting each other species (30% divided among 7 species). Interestingly, $\phi_k$ is relatively insensitive to $K$ ($\phi_k \simeq 0.32$ and 0.30 for $K=4$ and 16 respectively), but sensitive to the effective repertoire size ($\phi_k \simeq 0.19$ and 0.45 for a Simpson diversity of 1.5 and 3 respectively). For simplicity, in subsequent numerical simulations, we will assume that the sampling weight vector is the same as the switching weight vector.

Using the rescaled variables, the dynamic equations are

$$\begin{aligned}
\frac{ds_k}{dt} &= \mu\eta_k+\gamma y_k-s_k\left[(1-\phi_k)\beta_k c_k+\phi_k\sum_{j\neq k}\frac{v_j}{1-v_k}\beta_j c_j\right]-(\mu+\sigma)s_k+\sigma\sum_{j\neq k}a_{k,j}s_j,\\
\frac{dy_k}{dt} &= s_k\left[(1-\phi_k)\beta_k c_k+\phi_k\sum_{j\neq k}\frac{v_j}{1-v_k}\beta_j c_j\right]-(\mu+\nu+\gamma+\sigma)y_k+\sigma\sum_{j\neq k}a_{k,j}y_j,\\
\frac{dc_k}{dt} &= \tilde{\alpha}_k(1-c_k)\left[(1-\phi_k)y_k+\sum_{j\neq k}\phi_j\frac{v_k}{1-v_j}y_j\right]-\zeta_k c_k.
\end{aligned} \tag{S11}$$

##### S3-3.1 $R_0$ in the slow and fast switching limits

###### S3-3.1.1 The slow switching limit $\sigma \to 0$

Again, we will derive the flower-to-flower NGM by considering the dynamics near the disease-free equilibrium. Over its contaminated duration of $1/\zeta_k$, a contaminated species-$k$ flower will directly infect $\eta_k(1-\phi_k)\beta_k/\zeta_k$ species-$k$ specialists (when they forage on their specialty), and $\eta_i\phi_i\frac{v_k}{1-v_i}\beta_k/\zeta_k$ species-$i$ specialists for each $i \neq k$ (when they are randomly sampling). In turn, over its infectious lifetime of $\tau$, each infected species-$k$ specialist will directly contaminate $\tau\tilde{\alpha}_k(1-\phi_k)$ species-$k$ flowers and $\tau\tilde{\alpha}_j\phi_k\frac{v_j}{1-v_k}$ species-$j$ flowers for each $j \neq k$. Likewise, each infected species-$i$ specialist will directly contaminate $\tau\tilde{\alpha}_k\phi_i\frac{v_k}{1-v_i}$ species-$k$ flowers, $\tau\tilde{\alpha}_i(1-\phi_i)$ species-$i$ flowers, and $\tau\tilde{\alpha}_j\phi_i\frac{v_j}{1-v_i}$ species-$j$ flowers for each $j \notin \{i,k\}$. Altogether, the flower-to-flower NGM has matrix elements

$$\mathbf{Z}(0)_{j,k} = \frac{\tau\beta_k\tilde{\alpha}_j}{\zeta_k}\begin{cases} \eta_k(1-\phi_k)^2 + \sum_{i\neq k}\eta_i\phi_i^2\frac{v_k^2}{(1-v_i)^2}, & j=k, \\ \eta_k(1-\phi_k)\phi_k\frac{v_j}{1-v_k} + \eta_j(1-\phi_j)\phi_j\frac{v_k}{1-v_j} + \sum_{i\notin\{j,k\}}\eta_i\phi_i^2\frac{v_k v_j}{(1-v_i)^2}, & j\neq k. \end{cases} \tag{S12}$$

We see that unlike the single-major model, the flower-to-flower NGM is no longer diagonal. The single-species subnetworks are no longer disconnected, and the strength of connection is approximately linear in $\phi_k$. The larger the effective repertoire size, the larger the value of $\phi_k$, and so the stronger the between-subnetwork connections.

###### S3-3.1.2 The fast switching limit $\sigma \to \infty$

Just like in the single-major model, in the limit $\sigma \to \infty$, there is only one type of bee, a generalist. However, the fraction of time the generalist spends on flower species $k$ is no longer $\eta_k$, but instead given by

$$\eta_k(1-\phi_k) + \sum_{j\neq k}\eta_j\phi_j\frac{v_k}{1-v_j}. \tag{S13}$$

(One can verify that these fractions sum across $k$ to one.)

The derivation of $R_0$ follows the same argument as in Sec. S3-2.2.2 for the single-major model, except with $\eta_j$ replaced by the above fraction, so we have

$$R_{0,r}(\infty) = \tau\sum_k\left(\eta_k(1-\phi_k) + \sum_{j\neq k}\eta_j\phi_j\frac{v_k}{1-v_j}\right)^2\tilde{\alpha}_k\beta_k/\zeta_k, \tag{S14}$$

the subscript $r$ indicating that this formula applies to the random sampling model.

**An important note:** Although Eqn. (S14) looks different from its counterpart Eqn. (S9) for the single-major model, it really depends on how we parametrise the model. Our goal is to assess the consequences of ignoring floral constancy, so if the models are fitted to foraging data at the bee species level (which gives the fraction of time the bee species spends on each flower species), then all three models (single-major, random sampling, and double-major) should approach the same $\sigma \to \infty$ "individual generalist" limit and give the same numerical value of $R_0(\infty)$. Hence, the values of $\eta_k$ in Eqns. (S9) and (S14) should differ in such a way that the equations still give the same numerical results. Only when $\sigma$ is finite should the different notions of floral constancy between the three models start to manifest and result in different numerical values of $R_0(\sigma)$. On the other hand, if we have data at the individual bee level that allow us to parametrise $\eta_k$ directly, then $R_0(\infty)$ can vary between models.

###### S3-3.1.3 Numerical checks

Figure S3-5 shows a numerical test that confirms the fast- and slow-switching calculations of $R_0$ with $K = 8$ flower species, and randomly generated parameter sets. Prevalence was computed as the fraction of infected bees at time $t = 250$ for $\sigma = 25$ and $\sigma = 0$ respectively, solving the differential equations (S11) from initial conditions with disease prevalence 0.01 in all flower species at $t = 0$ and all bees susceptible at steady-state repertoire frequencies $\eta_{\{j,k\}}$. As expected, the behavior transitions from disease fade-out to disease persisteance at $R_0 = 1$ (indicated by the dashed vertical lines).

#### S3-4 The double-major model

In the double-major model, each bee has a repertoire comprising two different flower species, and spends half of its time foraging on each of them. Repertoires are labelled by unordered sets $\{j,k\}$ with $j \neq k$, so the state variables $S_k$ and $I_k$ in the single-major model are replaced by $S_{\{j,k\}}$ and $I_{\{j,k\}}$. We assume that any pair of species is allowed as a repertoire, so there are $\binom{K}{2} = K(K-1)/2$ possible repertoires.

We assume that each bee switches repertoire by changing one flower species at a time, with equal probability of dropping either species $j$ or $k$ in the current repertoire $\{j,k\}$. Having dropped a species, the bee gains species $m \notin \{j,k\}$ as its replacement with probability $a_{m|\{j,k\}}$. Therefore, $\{j,k\}$ switches to $\{m,k\}$ or to $\{m,j\}$ each at the same rate of $(\sigma/2)a_{m|\{j,k\}}$. At the disease-free steady state, $\eta_{\{j,k\}}$ gives the fraction of bees with repertoire $\{j,k\}$, and again we assume that new bees are recruited in the same fractions.

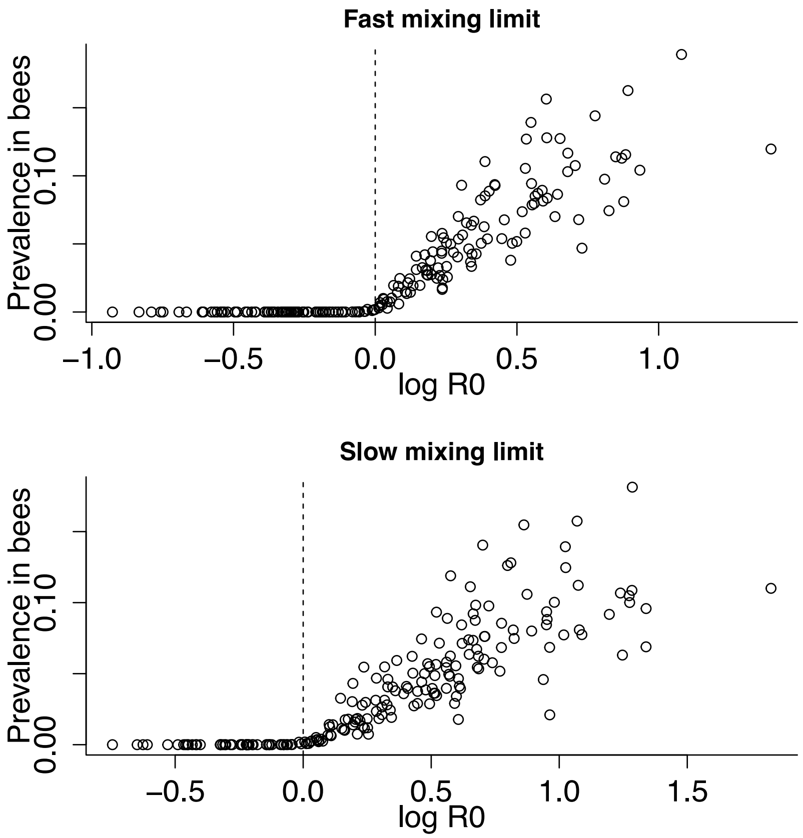

Figure S3-5: Numerical test of the calculations for $R_0$ in the fast- and slow-switching limits of the random sampling model using randomly generated parameter sets. See text for details. Figure made by R script `test_R0_random_sampling.R` and scripts that it sources.

Except for $S_k$, $I_k$, $a_{k,j}$ and $\eta_k$ (which are replaced by their double-major analogues described above), all other state variables and parameters carry over from the original model. Using rescaled variables, the dynamic equations are

$$\begin{aligned}
\frac{ds_{\{j,k\}}}{dt} =& b\eta_{\{j,k\}} + \gamma y_{\{j,k\}} - 0.5(\beta_k c_k + \beta_j c_j)s_{\{j,k\}} - (\mu+\sigma)s_{\{j,k\}} \\
&+ \frac{\sigma}{2}\sum_{m\notin\{j,k\}} a_{j|\{m,k\}} s_{\{m,k\}} + \frac{\sigma}{2}\sum_{m\notin\{j,k\}} a_{k|\{m,j\}} s_{\{m,j\}} \\
\frac{dy_{\{j,k\}}}{dt} =& 0.5(\beta_k c_k + \beta_j c_j)s_{\{j,k\}} - (\mu+\nu+\gamma+\sigma)y_{\{j,k\}} \\
&+ \frac{\sigma}{2}\sum_{m\notin\{j,k\}} a_{j|\{m,k\}} y_{\{m,k\}} + \frac{\sigma}{2}\sum_{m\notin\{j,k\}} a_{k|\{m,j\}} y_{\{m,j\}} \\
\frac{dc_k}{dt} =& 0.5\tilde{\alpha}_k(1-c_k)\sum_{m\neq k} y_{\{m,k\}} - \zeta_k c_k,
\end{aligned} \tag{S15}$$

where

$$s_{\{j,k\}} \equiv S_{\{j,k\}}/M, \quad y_{\{j,k\}} \equiv I_{\{j,k\}}/M. \tag{S16}$$

Again the specialty fraction $\eta_{\{j,k\}}$ is a derived parameter that can be related to other model parameters. If we further assume that the switching probabilities can be parametrised by a switching weight vector $\{w_1, w_2, ..., w_K\}$ of unit sum such that

$$a_{m|\{j,k\}} = \frac{w_m}{1-w_j-w_k}, \tag{S17}$$

then one can show that

$$\eta_{\{j,k\}} = \frac{w_j w_k (1-w_k-w_j)}{Z}, \quad Z = \sum_{r<s} w_r w_s (1-w_r-w_s). \tag{S18}$$

The demonstration of (S18) consists of confirming that the $\eta_{\{j,k\}}$ satisfy detailed balance, i.e. the probability flow from one repertoire to another is equal to the flow in the opposite direction between those two repertoires, when the state frequencies are (S18). The steady-state flow from $S_{\{j,k\}}$ to $S_{\{m,k\}}$ where $m \notin \{j,k\}$ is

$$\frac{\sigma}{2}\eta_{\{j,k\}} a_{m|\{j,k\}} = \frac{\sigma}{2}(w_j w_k (1-w_k-w_j)) \left( \frac{w_m}{1-w_j-w_k} \right) / Z = \frac{\sigma}{2} w_j w_k w_m / Z.$$

As $j$ and $m$ are arbitrary, the flow in the opposite direction is the same, as claimed.

#### S3-4.1 $R_0$ in the slow and fast switching limits

##### S3-4.1.1 The slow switching limit $\sigma \to 0$

In this limit, bees are born with some repertoire and are assumed to stick with it for life, with $\eta_{\{j,k\}}$ the fraction having repertoire $\{j,k\}$.

An NGM that includes infected bees must somehow reduce the two-dimensional classification by repertoire into a one-dimensional state vector. To avoid that, we choose to derive the $K \times K$ flower-to-flower NGM. To compute $\mathbf{Z}(0)_{i,j}$ we consider starting the rescaled model (S15) with a very low level of infection in flower species $j$, $c_j(0) = \varepsilon$, and all bees susceptible so $s_{\{i,j\}}(0) = \eta_{\{i,j\}}$. Infection passes in one "generation" from flower species $j$ to flower species $i \neq j$ only through bees with repertoire $\{i,j\}$. The initially infected species $j$ flowers remain infected for average time $1/\zeta_j$, producing infection level $y_{\{i,j\}} = 0.5\varepsilon\beta_j\eta_{\{i,j\}}/\zeta_j$ among bees with repertoire $\{i,j\}$. Each of those bees remains infected for time $\tau$ on average, giving rise each to $0.5\tau\tilde{\alpha}_i$ contaminated flowers in species $i$. We therefore have

$$\mathbf{Z}(0)_{i,j} = 0.25\tau\tilde{\alpha}_i\beta_j\eta_{\{i,j\}}/\zeta_j, \quad i \neq j. \tag{S19}$$

Infection passes from flower species $j$ back to itself (i.e., back to other individuals of the same species) through bees that have any repertoire $\{k,j\}, k \neq j$. Applying the same arguments as above to each of these repertoires, we have

$$\mathbf{Z}(0)_{j,j} = 0.25\tau\tilde{\alpha}_j\beta_j\sum_{k\neq j}\eta_{\{k,j\}}/\zeta_j = 0.25\tau\tilde{\alpha}_j\beta_j\rho_j/\zeta_j, \tag{S20}$$

where $\rho_j$ is the fraction of bees that have flower species $j$ in their repertoire, i.e.

$$\rho_j = \sum_{k\neq j}\eta_{\{k,j\}}. \tag{S21}$$

$R_0(0)$ is then, as usual, the dominant eigenvalue of $\mathbf{Z}(0)$.

Like the random sampling model, the flower-to-flower NGM is no longer diagonal and so the single-species subnetworks are no longer disconnected. Nonetheless, we will show in Sec. S3-4.1.3 that the connections between subnetworks are still weaker than in the case of no floral constancy.

###### S3-4.1.2 The fast switching limit $\sigma \to \infty$

In the limit of $\sigma \to \infty$, again there is only one kind of bee, which spends a fraction $\eta_{\{i,j\}}$ of its time having repertoire $\{i,j\}$. Hence, the derivation of the one-dimensional "bee-to-bee" NGM follows the same argument as in the Ellner et al. (2020) single-major model.

To recap, once infected, a bee spends average time $\tau$ until it recovers, and therefore average total time $0.5\tau\rho_j$ infecting flower species $j$. This produces $0.5\tau\rho_j\tilde{\alpha}_j$ contaminations in flower species $j$ in the rescaled model. Each such contaminated flower remains contaminated for average duration $1/\zeta_j$, resulting in a time-integrated integrated total force of infection of $0.5\tau\rho_j\tilde{\alpha}_j\beta_j/\zeta_j$ from all first-generation contaminated species $j$ flowers. Uninfected bees there are exposed to this force of infection only while they have species $j$ in their repertoire (and thus spend half their time foraging on species $j$), so the resulting number of second-generation bee infections is $0.25\tau\rho_j^2\tilde{\alpha}_j\beta_j/\zeta_j$. Overall bee-to-flower-to-bee transmission is the sum of these, giving

$$R_{0,d}(\infty) = 0.25\tau\sum_{j=1}^{K}\rho_j^2\tilde{\alpha}_j\beta_j/\zeta_j \tag{S22}$$

the subscript $d$ indicating that this formula applies to the double-major model.

Compared to Eqn. (S9), here we have replaced $\eta_j$ (the fraction of bees with species $j$ as their specialty) by $0.5\rho_j$ (where $\rho_j$ is the fraction of bees with species $j$ in their repertoire). The definitions of $\eta_j$ and $\rho_j$ are similar, but we note that $\sum_j \eta_j = 1$, whereas $\sum_j \rho_j = 2$ because each bee has two flowers in its repertoire. Thus, $\rho_j/2$ has the same biological meaning in this model that $\eta_j$ has in the single-major model: it is the fraction of total bee foraging time (by all bees) that bees spend foraging at flower species $j$, at steady state and in the rapid switching limit, in the absence of disease.

###### S3-4.1.3 Further analysis

While we have already derived the bee-to-bee NGM in the fast switching ($\sigma \to \infty$) limit, it is informative to also derive the flower-to-flower NGM and compare with the slow switching ($\sigma \to 0$) limit. In the $\sigma \to \infty$ limit, the flower-to-flower NGM has matrix elements

$$\mathbf{Z}(\infty)_{i,j} = 0.25\tau\tilde{\alpha}_i\beta_j\rho_i\rho_j/\zeta_j \qquad \text{(S23)}$$

for all $i$ and $j$. Clearly $\mathbf{Z}(0)_{j,j} > \mathbf{Z}(\infty)_{j,j}$. For $i \neq j$, we conjecture that $\eta_{\{i,j\}} < \rho_i\rho_j$ (this is unproven but supported by simulations using randomly generated switching weights), so we also have $\mathbf{Z}(0)_{\{i,j\}} < \mathbf{Z}(\infty)_{\{i,j\}}$.

Hence, we see that in the slow switching limit, within-species pathogen flow is still stronger and between-species flow still weaker than in the fast switching limit. This is because in the slow switching limit, bees are more likely to re-visit the same species of flowers they were infected from, simply by virtue of the fact that the species they were infected from must have been in their repertoires in the first place.

###### S3-4.1.4 Numerical checks

Figure S3-6 shows a numerical test that confirms the fast- and slow-switching calculations of $R_0$ with $K = 4$ flower species, and randomly generated parameter sets. The test setup is the same as in Sec. S3-3.1.3. As expected, the behavior transitions from disease fade-out to disease persisteance at $R_0 = 1$ (indicated by the dashed vertical lines).

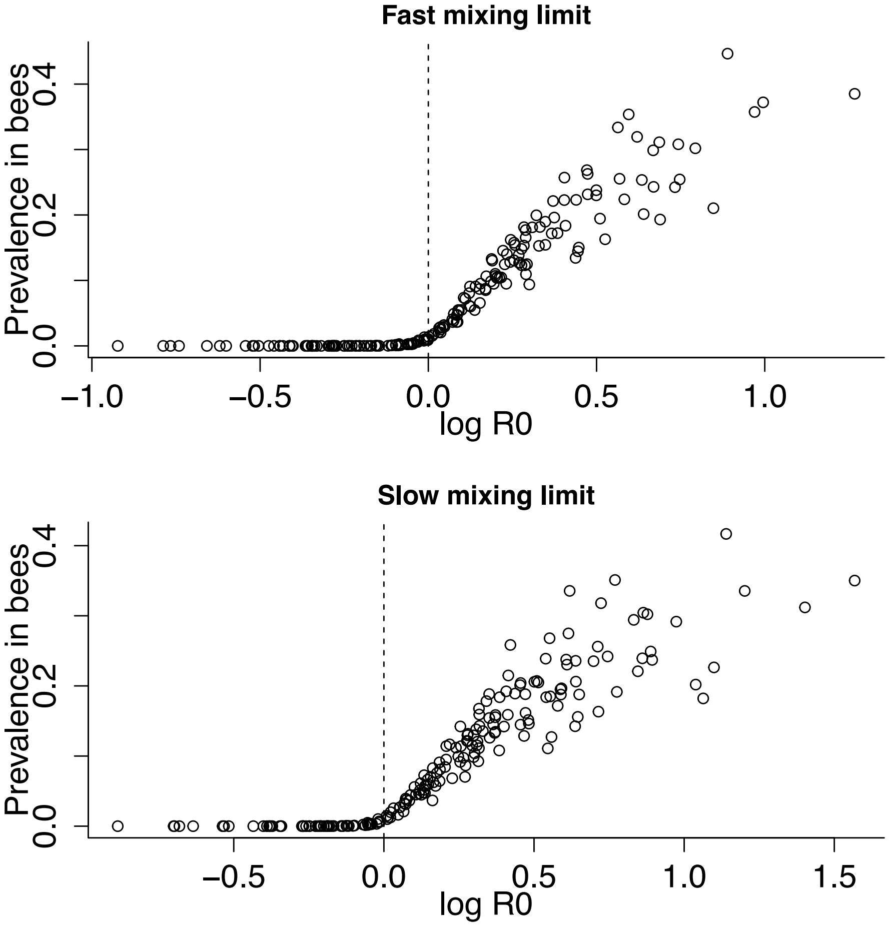

Figure S3-6: Numerical test of the calculations for $R_0$ in the fast- and slow-switching limits of the double-major model using randomly generated parameter sets. See text for details. Figure made by R script `test_R0_double_major.R` and scripts that it sources.

Eguíluz, V. M., and K. Klemm. 2002. Epidemic threshold in structured scale-free networks. Phys. Rev. Lett. 89:108701.

Ellner, S. P., W. H. Ng, and C. R. Myers. 2020. Individual specialization and multihost epidemics: Disease spread in plant-pollinator networks. American Naturalist 195:E118–E131.

Heinrich, B. 1976. The foraging specializations of individual bumblebees. Ecological Monographs 46:105 – 128.

Truitt, L. L., S. H. McArt, A. H. Vaughn, and S. P. Ellner. 2019. Trait-based modeling of multi-host pathogen transmission: Plant-pollinator networks. American Naturalist 193:E149–E167.
